# Evaluation of T Cell Receptor Construction Methods from scRNA-Seq Data

**DOI:** 10.1101/2024.11.01.621447

**Authors:** Ruonan Tian, Zhejian Yu, Ziwei Xue, Jiaxin Wu, Lize Wu, Shuo Cai, Bing Gao, Bing He, Yu Zhao, Jianhua Yao, Linrong Lu, Wanlu Liu

## Abstract

T cell receptors (TCRs) serve pivotal roles in the adaptive immune system by enabling recognition and response to pathogens and irregular cells. Various methods exist for TCR construction from single-cell RNA sequencing (scRNA-seq) datasets, each with its unique characteristics regarding accuracy, sensitivity, adaptability, usability, time, and memory consumption. Yet, a comprehensive understanding of their relative strengths and weaknesses for different applications remains elusive. In our research, we implemented a benchmark analysis utilizing experimental single-cell immune profiling datasets encompassing paired scRNA-seq as input and scTCR-seq datasets as ground truth reference from human and mouse. Additionally, we introduced a novel simulator, YASIM-scTCR (Yet Another Simulator for single-cell TCR), capable of generating scTCR-seq reads containing a diverse array of TCR-derived sequences under different sequencing depths and read lengths. Our results consistently showed that TRUST4 outperformed others across multiple datasets, while MiXCR and DeRR also demonstrated considerable accuracy. We also discovered that the sequencing depth inherently imposes a critical constraint on successful TCR construction from scRNA-seq data. In summary, we present a benchmark study to aid researchers in choosing the most appropriate methods for reconstructing TCR from scRNA-seq data.

## INTRODUCTION

T cell receptors (TCRs) play central roles in recognizing pathogen- and self-derived antigens, thereby fostering immunosurveillance of infectious diseases, autoimmune disorders, and cancers [1,2]. TCRs are highly diverse due to V(D)J recombination and the pairing of α/β chains, especially in the antigen-recognizing complementary-determining region 3 (CDR3), leading to an enormous TCR repertoire capable of recognizing a wide range of antigens [3,4]. Therefore, characterizing the antigen receptor repertoire and understanding its dynamics in disease progression is crucial for guiding vaccine design and developing precise immunotherapy [5,6].

Next-generation sequencing (NGS)-based bulk TCR-seq enables the characterizing of antigen receptor repertoires under different disease conditions [7–9]. However, the lack of paired information for α/β chains limits its wide application in downstream experimental validation. Emerging single-cell technologies offer a promising approach for capturing gene expression profiles (GEX) alongside paired TCR α/β chains information at the single-cell level, referred to as single-cell immune profiling (scRNA+TCR-seq) [9–11]. This approach potentially provides profound insights into antigen receptor repertoire analysis and helps reveal functional TCRs [9,10]. While single-cell immune profiling data based on 10X and SMART-seq methodologies has seen exponential growth, with the 10X dataset being the primary strategy, its widespread application is still restrained due to the high cost and additional TCR amplification steps required in library construction [9,12,13]. For instance, in our latest version of the single-cell immune profiling-based database huARdb (human Antigen Receptor database), we only collect around half a million T cells [14], significantly fewer than the theoretically predicted TCR diversity of 10^20^ [15]. Meanwhile, scRNA-seq datasets of T cells, inherently containing TCR sequence information, are abundantly accessible in public databases [11,16]. Nevertheless, TCR-derived sequences are frequently disregarded in most scRNA-seq studies as they might not constitute the primary focus of investigations. However, it is worth noting that even with lower sensitivity, scRNA-seq may still offer researchers the potential to reconstruct TCR sequence information from the scRNA-seq data, which could significantly enhance our comprehensive understanding of the functions of T cells [17,18].

TCR construction methods share certain similarities with *de novo* transcriptome assembly and contig annotation, such as Trinity [12] and IgBLAST [19]. However, it also diverges from these general-purpose assemblers in several key aspects. Firstly, there is a lack of reliable reference for the highly variable CDR3 regions of TCR [3]. Secondly, there is a need for a specific reference for VJ genes rather than a genome-wide reference [10]. These distinct characteristics render TCR assembly and annotation more challenging compared to general-purpose assemblers, highlighting the need for specialized methods [10].

Various steps are involved in TCR construction, including candidate reads identification from bulk RNA-seq or scRNA-seq ((sc)RNA-seq), TCR reads assembly, and V(D)J gene annotation. While the recombination processes of TCRs and B-cell receptors (BCRs) share significant similarities, they also possess distinct differences. Therefore, to facilitate a comprehensive summary of method characteristics, we conducted a thorough review of all available methods capable of reconstructing TCRs and BCRs from (sc)RNA-seq datasets, including MiXCR [20], TraCeR [21], VDJer [22], BASIC [23], BALDR [24], BraCeR [25], VDJPuzzle [26], ImRep [27], CATT [28], TRUST4 [29] and DeRR (Supplementary Table 1). While BALDR, BraCeR, and VDJer exclusively support BCR construction, TraCeR and DeRR are specifically designed for TCR construction. Generally, these methods can be categorized based on whether they implemented a *de novo* algorithm in TCR/BCR construction. For example, MiXCR, VDJer, ImRep, CATT, and TRUST4 adopt a *de novo* approach for read alignment, assembly, and annotation in TCR/BCR construction. Other methods, such as TraCer, BASIC, BLADR, BraCer, VDJPuzzle, and DeRR construct a pipeline by incorporating existing alignment, assembly, or annotation methods for such task. Commonly used methods for read alignment in such pipelines include Bowtie [30], Bowtie2 [31], BWA [32], STAR [33], while Trinity [12] is often employed for assembly, and IgBLAST [19] is adapted for VJ gene and CDR3 amino acid sequence annotation. In comparison to *de novo* algorithms, methods wrapping several existing packages may be less user-friendly due to efforts in satisfying their prerequisites.

To provide a comprehensive resource for researchers to select suitable TCR construction methods according to their unique needs, we conducted an extensive evaluation of the performance of current TCR construction methods on scRNA-seq, scTCR-seq, pseudo-bulk RNA-seq, bulk TCR-seq, and simulated scTCR-seq data in this study. The seven methods applications included in this benchmark study were MiXCR, TraCeR, BASIC, ImRep, CATT, TRUST4, and DeRR. Leveraging previously published scRNA+TCR-seq datasets, we utilized scRNA-seq data as input for various methods and employed scTCR-seq dataset as ground truth to assess methods accuracy and sensitivity, focusing on CDR3 amino acid sequences and VJ gene usages. Additionally, we generated pseudo bulk RNA-seq data to examine the effect of cell number on TCR construction accuracy. Furthermore, we developed a simulator called YASIM-scTCR (Yet Another SIMulator for single-cell TCR) to generate scTCR-seq reads containing TCR- and non-TCR-derived sequences, thus allowing us to evaluate methods performance under various sequencing depths and read lengths. This analysis with simulated data facilitated the identification of potential algorithmic limitations, thereby fostering the development of improved methods. Finally, we also assessed the computational efficiency of different methods. In conclusion, our analyses resulted in the establishment of quality control metrics for the precise assembly of TCRs from scRNA-seq data.

## RESULTS

### Overview of Methods Supporting TCR Reconstruction from scRNA-seq Data

To evaluate the performance of various TCR construction methods, we compiled previously published single-cell immune profiling datasets containing paired scRNA-seq and scTCR-seq data from both human and mouse (Supplementary Table 2). By subjecting 10X scRNA-seq data to different TCR construction methods and comparing the assembled TCR with its paired 10X scTCR-seq data, we thoroughly assessed the accuracy and sensitivity of each method (Fig. 1A). In general, the TCR construction algorithms can be categorized into three main steps: The first step searched for candidate TCR-derived reads from FASTQ files using reference V(D)J segment sequences or BAM files from aligned genomic coordinates; They are then assembled to TCR contigs using methods like de Bruijn Graph; Lastly, V(D)J segments are identified and sequences of CDR3 regions are annotated (Fig. 1A).

**Figure 1.**
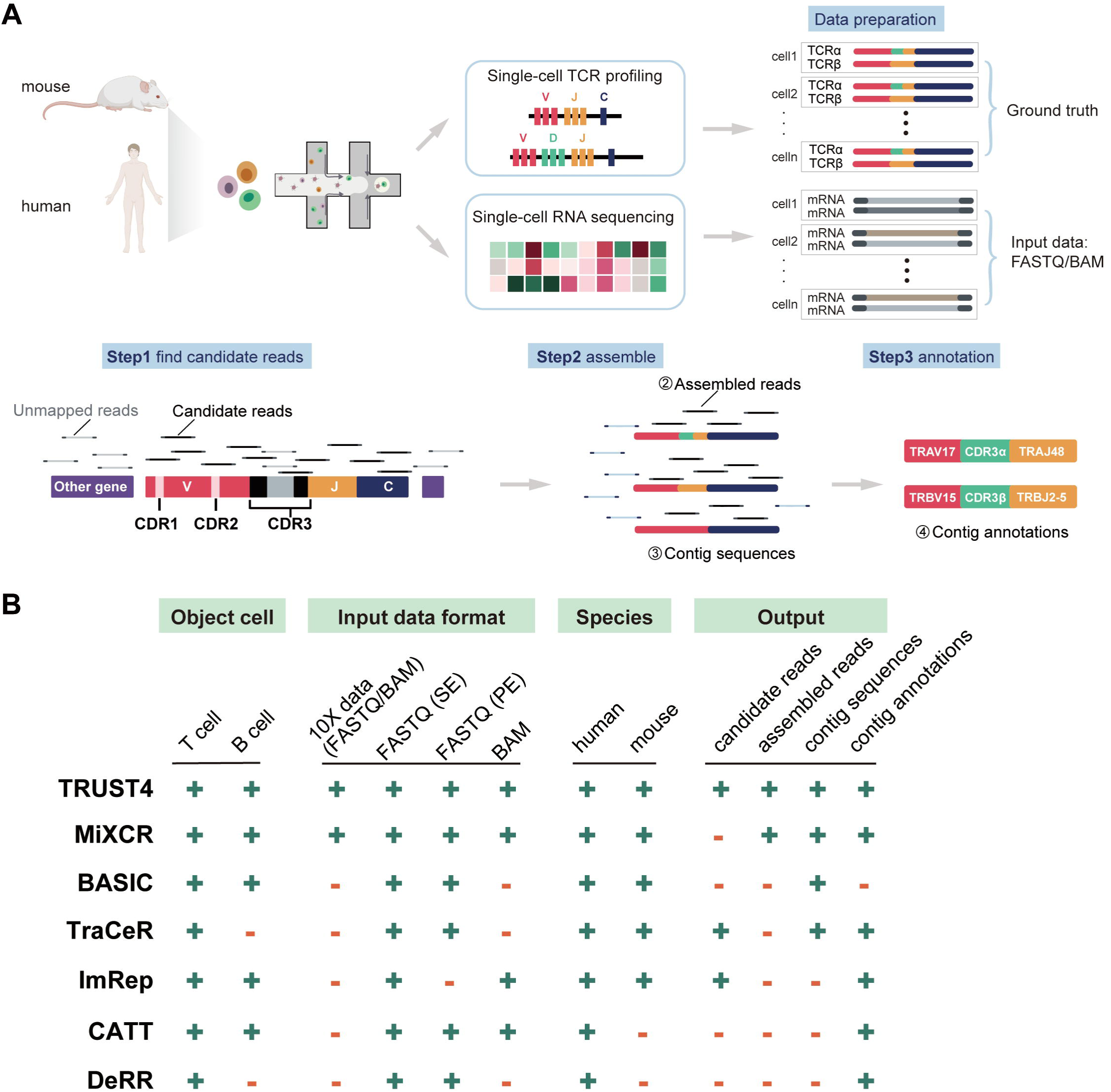
A benchmark framework for TCR reconstruction methods using real single-cell RNA-seq data. **A.** Schematic representation of the benchmark workflow. Human and mouse scRNA-seq and scTCR-seq datasets were used as input data and ground truth for downstream analyses. The analytical pipeline comprised three key steps: (i) identifying candidate reads from the input data, (ii) assembling contigs from all candidate reads with partial overlaps, and (iii) annotating the contig sequences and reconstructing TCRs. BAM, Binary Alignment Map format. **B.** Comprehensive evaluation of the adaptability of various methods with a focus on four key aspects: object cell, input data format, species, and output. “+” represents the presence of this feature, and “-” represents the absence of this feature.

In this benchmark study, we comprehensively analyzed the performance of seven distinct methods designed for TCR construction from (sc)RNA-seq data. Among these, six have been previously published in peer-reviewed literature [20,21,23,27–29], while DeRR is publicly available [34]. We first summarized the adaptability of these methods across four different aspects: object cell type (T/B cell), input data format (FASTQ/BAM files), species supported (human or mouse), and output schema (Fig. 1B). Regarding the input data format, all methods supported raw FASTQ format, and a subset of them (TRUST4, MiXCR, ImRep, and CATT) also accommodated BAM format. While all methods are compatible with demultiplexed scRNA-seq data as individual cells, such as SMART-seq, it is worth noting that currently only MiXCR and TRUST4 support 10X scRNA-seq (Fig. 1B). Furthermore, we summarized the output schema, which encompassed candidate read sequences, assembled read sequences, contig sequences, and contig annotations, as these elements play a crucial role in conducting for comprehensive benchmarking analyses (Fig. 1B). Our in-depth evaluation revealed considerably variability among these methods in terms of the information provided in their outputs (Fig. 1B). For instance, TRUST4 demonstrated a commendable capability in offering comprehensive output at each stage of the analysis process, while BASIC only provided the contig sequences without any annotation, leading us to exclude it in the accuracy and sensitivity analysis (Fig. 1B).

### Accuracy and Sensitivity of Different Methods Using Experimental scRNA-seq Data

To investigate the impact of input file format, we evaluated various methods using both FASTQ and genome-aligned BAM files for scRNA-seq data (Fig. 2). We assessed the accuracy and sensitivity of CDR3 amino acid sequences, J/V gene calling and Assembled-TCR (AsTCR) for both α and β chains, using 10X scRNA-seq data from human and mouse in both format (Fig. 2 and Supplementary Table 2). The results revealed that TRUST4 (FASTQ or BAM) and MiXCR (BAM) demonstrated consistently highest sensitivity in assembling both α and β chains (Fig. 2A). MiXCR(FASTQ), CATT (FASTQ), DeRR, and TraCeR exhibited commendable performance in CDR3, J gene, or V gene calling, while CATT (BAM) showed comparatively lower sensitivity in this context (Fig. 2A). ImRep (FASTQ and BAM) also exhibited comparable performance in assembly of CDR3 regions on either chain. However, it only reported J genes without a specific subgroup for the β chains leading to uncharacterized TRBJ gene calling (Fig. 2A). Regarding accuracy, the performance of methods in CDR3, J gene, and V gene calling was generally consistent, with DeRR and MiXCR displayed the highest accuracy on AsTCR (Fig. 2B).

**Figure 2.**
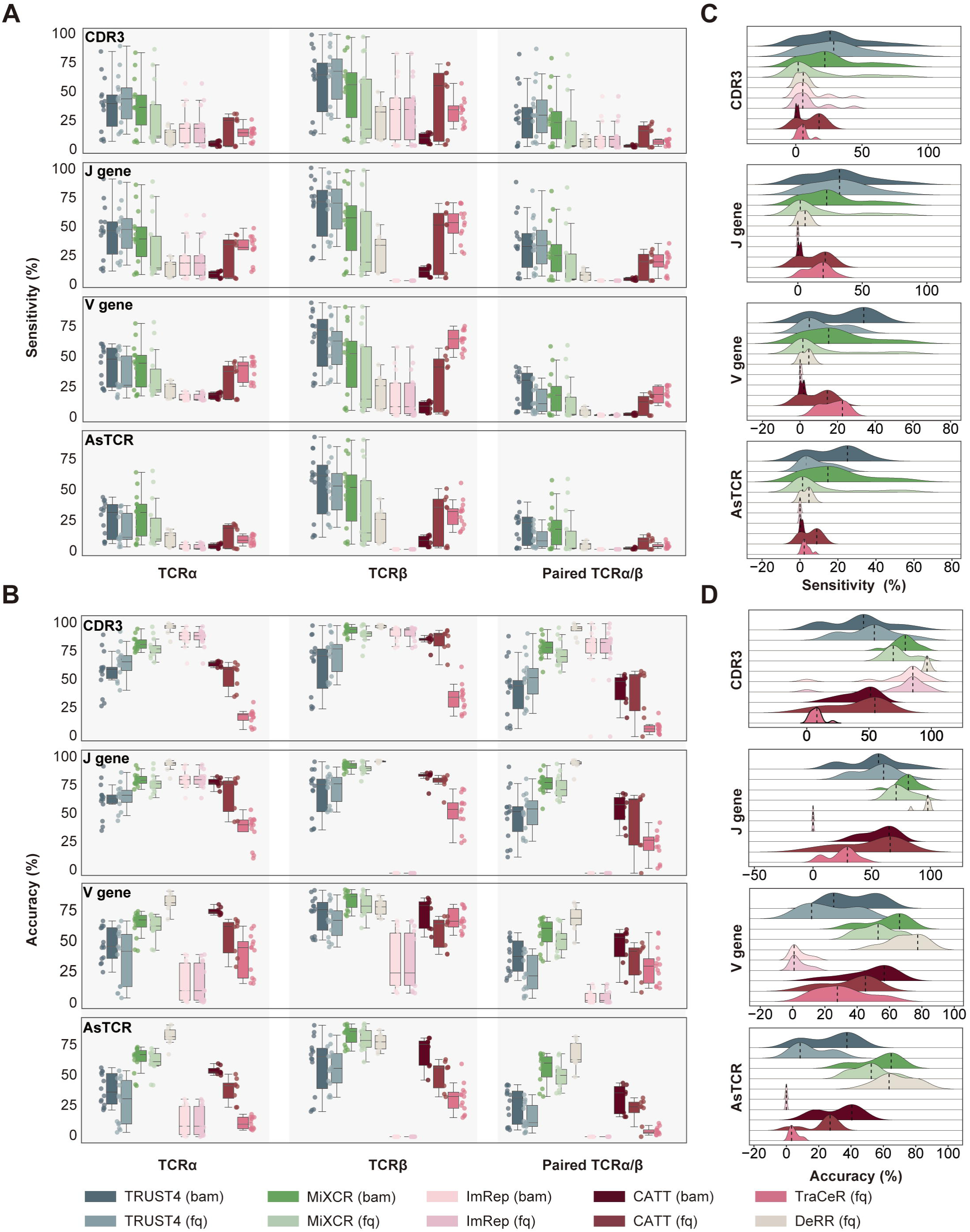
Comparative analysis of TCR construction methods. **A.** Box plot represents the sensitivity of TCR prediction of different methods. The line in the middle represents the median; boxes represent the 25th (bottom) and 75th (top) percentiles; and whiskers represent the minimum and maximum points within 1.5 times the interquartile range. **B.** Box plot represents the accuracy of TCR prediction of different methods. **C.** Ridge plot represents the sensitivity distribution of CDR3, V gene, J gene, and AsTCR of paired TCRα/β. **D.** Ridge plot represents the accuracy distribution of CDR3, V gene, J gene, and AsTCR of paired TCRα/β. CDR3, complementarity-determining region 3; AsTCR, assembled TCR; fq, FASTQ format; bam, Binary Alignment Map format.

For both accuracy and sensitivity evaluation, our results revealed that TCRβ chains overall outperformed TCRα chains across all evaluated methods (Fig. 2A, B). This observation could be attributed to several factors. Firstly, TCRα chains typically display lower expression levels compared to TCRβ chains (Supplementary Fig. 1A). Furthermore, the TCRβ chain often receives more attention from researchers, potentially leading to a preference towards the TCRβ chain in the design of TCR construction algorithms and reference sequences. We also observed variations in the performance of TCR construction when using FASTQ inputs compared to BAM format files (Fig. 2A, B). This discrepancy could be potentially attributed to the alignment process, where candidate sequences with a higher mutation rate may fail to align correctly and thus get discarded. In summary, TRUST4 and MiXCR demonstrated notably higher sensitivity for TCR construction, while DeRR and MiXCR exhibited relatively higher accuracy levels.

The scRNA-seq data generated by the 10X Genomics platform serves as a typical example of droplet-based library construction technology. This platform is capable of producing paired scRNA-seq and scTCR-seq data, thus we utilized the paired scTCR-seq as ground truth for TCR construction from scRNA-seq. In addition, we scrutinized the TCR assembly performance using 50 cells generated by the SMART-seq library preparation (Supplementary Table 2). However, given the lack of paired scTCR-seq data for SMART-seq, we are unable to formulate the ground truth. To assess the performance of various methods, we quantified the number of cells with AsTCRs and calculated their sharing (Supplementary Fig. 2A-D). TRUST4 assembled the most AsTCRs, and high overlaps for V/J genes from TCRβ chains with other methods, while TCRα chains displayed minimal consistency among those tested methods (Supplementary Fig. 2A-D).

During the accuracy and sensitivity analysis, we observed substantial inconsistency for these TCR construction methods when applied to different scRNA-seq datasets (Fig. 2C, D). Additionally, all evaluated methods exhibited limited sensitivity in TCR construction, with the best-performing method, TRUST4 (BAM), achieving an average sensitivity of only 25% for assembled TCRα/β chains (Fig. 2A). These findings prompted us to investigate the underlying factors. Specifically, we would like to address two questions: 1) How do variations in scRNA-seq data itself, such as species origin, sequencing strategies (paired-end or single-end), sequencing length, and sequencing depth, impact the performance of these methods? 2) Is the limitation associated with the algorithms themselves, or could it be attributed to the lack of capturing for TCR sequences? For example, without TCR-specific amplification, most scRNA-seq data may not capture enough TCR sequences, thereby hindering the construction of TCRs in certain cells.

### Accuracy and Sensitivity of Different Methods Using Simulated scTCR-seq Data

To address the aforementioned challenges, we developed a simulation framework called YASIM-scTCR based on realistic parameters estimated from single-cell immune profiling datasets of 1.08 million T cells [14,35]. Using YASIM-scTCR, we generated scTCR-seq data containing TCR-derived reads (Fig. 3A). In this simulation framework, YASIM-scTCR initially generates TCR contigs by adhering to the principles of TCR recombination (Fig. 3A). Specifically, V and J genes are selected from the human V and J gene reference annotations, and deletions are then introduced at the V-CDR3 (V gene-CDR3) and CDR3-J (CDR3-J gene) junctions. Subsequently, CDR3 sequences are generated and combined with the deleted V/J gene segments to form TCR contigs. To ensure the simulation of TCRs with realistic characteristics, we performed statistical analysis on 1.08 million public T cells single-cell immune profiling datasets [14,35], which encompassed the length distribution of full-length TCRα/β chains, as well as CDR3α/β regions (Supplementary Fig. 3A-C). To control the potential number of deleted amino acids at the V-CDR3 and CDR3-J junctions, we evaluated the frequency distribution for the number of deletions and the length distribution for J genes (Supplementary Fig. 3D, E). Additionally, we assessed and simulated amino acid preferences in the CDR3α/β (Supplementary Fig. 3F, G). We also evaluated and simulated the distribution of V/J gene usage bias (Supplementary Fig. 4). Subsequently, we constructed complete TCR contigs based on the V(D)J sequences, incorporating C gene and cell barcode information, in line with the 5’ 10X Genomics scTCR-seq library construction strategy (Fig. 3A). After generating the TCR contigs, YASIM-scTCR would simulate gene expression profiles for protein-coding genes based on experimental scRNA-seq data using scDesign2 [36]. Lastly, by employing ART [37], a tool for generating synthetic NGS reads, we complied the simulated scTCR-seq data containing TCR- and non-TCR-derived reads (Fig. 3A). Moreover, YASIM- scTCR offers flexibility in the simulation process by allowing the adjustment of various parameters, such as read length and sequencing depth over TCR.

**Figure 3.**
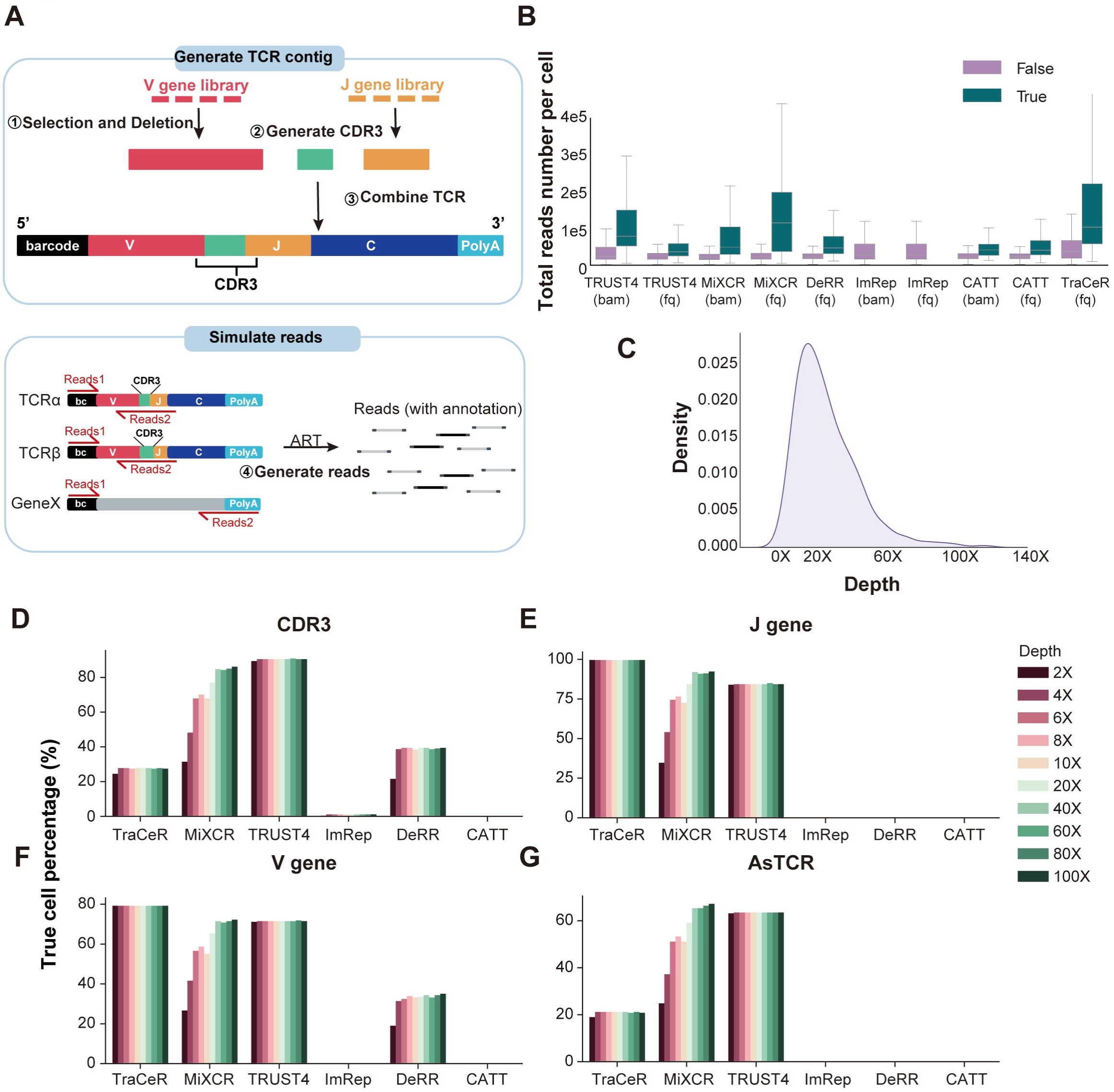
Performance comparison across real and simulated datasets with different sequencing depth. **A.** Schematic representation of simulating scTCR-seq and scRNA-seq datasets. bc, cell barcode; PolyA, a stretch of adenine-rich nucleotides added to the 3’ end of RNA molecules; Red box, V gene; Green box, CDR3 sequence; Orange box, J gene; Dark blue box, C gene; Light blue box, polyA sequence; Black box, cell barcode suquence; Gray box, non-TCR genes. **B.** Box plot represents the total read number of cells with successfully (True) and unsuccessfully (False) assembled TCR on the collected experimental scRNA-seq datasets. The line in the middle represents the median; boxes represent the 25th (bottom) and 75th (top) percentiles; and whiskers represent the minimum and maximum points within 1.5 times the interquartile range. **C.** Density plot represents the sequencing depth distribution of TCR sequences in TCR- assembled cells obtained from TRUST4 **D-G.** Bar plots represent the performance of paired TCRα/β reconstruction on the simulated datasets with different sequencing depth. CDR3, complementarity-determining region 3; AsTCR, assembled TCR; fq, FASTQ format; bam, Binary Alignment Map format.

Using YASIM-scTCR, we conducted an in-depth analysis of candidate reads and their impact on the performance of TCR construction. Among the methods examined, TRUST4, TraCeR, and ImRep were the ones capable of generating comprehensive outputs containing information on candidate reads. For the other methods such as DeRR, we made necessary modifications to its source code to enable the output of candidate reads. Subsequently, we assessed the efficiency of candidate read identification and the final TCR recombination accuracy using simulated scTCR-seq data encompassing TCR reads from 500 cells. In terms of candidate reads identification, TRUST4 and TraCeR displayed exceptional performance, achieving 100% for both true positive rates (TPR) and true negative rates (TNR) (Supplementary Fig. 5A). DeRR also reported a high level of true positive candidate rates (100% TPR), although it exhibited a minimal TNR (0%) (Supplementary Fig. 5A). Conversely, ImRep exhibited a high TNR (96%) along with a low TPR (68%), potentially influencing its overall TCR construction performance (Supplementary Fig. 5A). The analysis of TCR construction accuracy over simulated data corresponded closely with the TPR in candidate read identification (Supplementary Fig. 5B). Specifically, TRUST4 outperformed in CDR3, J gene, V gene calling for both α and β chains, while TraCeR achieved high accuracy in J gene and V gene calling (Supplementary Fig. 5B). DeRR achieved approximately 20-40% accuracy in AsTCR construction, whereas ImRep mostly obtained less than 5% accuracy in this task (Supplementary Fig. 5B). We also observed that all methods, particularly TRUST4, performed significantly better on the simulated data compared to experimental scRNA-seq data. Given that YASIM-scTCR simulated 400X scTCR-seq reads in this scenario, which may contain more TCR-derived reads compared to the experimental scRNA-seq data. This observation partially corroborates our hypothesis that the low TCR construction rate in scRNA-seq data may be primarily attributed to the lack of TCR information in the data itself rather than inherent deficiencies in the software algorithms.

### Performance Analysis for Different Methods across Various Species Origins, Sequencing Strategies, Read Lengths, and Depths

To elucidate potential factors contributing to the varied performance of TCR construction methods, we initially visualized the performance of these methods separately for human and mouse samples. While most TCR construction methods exhibited comparable accuracy and sensitivity levels for both species, notable heterogeneity within each species suggested additional factors may influence the performance of the methods (Supplementary Fig. 6A, B).

To assess the impact of scRNA-seq library construction strategies, we segregated performance of the methods based on the sequencing strategies either single-end and paired-end sequencing corresponding to the 10X Genomics Single Cell 3’ and 5’ Gene Expression (GEX) libraries, respectively (Supplementary Table 3-4). Our comparative analysis revealed that paired-end sequencing data demonstrated higher accuracy and sensitivity compared to single-end sequencing data (Supplementary Fig. 6C, D). This observation aligns with the approach of single-cell immune profiling techniques, suggesting that 3’ GEX library construction strategies may inadequately capture full-length TCR sequences. Consequently, we recommend the use of scRNA-seq data from the 5’ GEX Library for TCR construction.

Next, we investigated the influence of read length for TCR construction using simulated scTCR-seq data with varying read lengths, while maintaining a fixed sequencing depth of 400X. The tested read lengths included 100bp (base pair), 150bp, and 250bp (Supplementary Fig. 7). A read length of 150bp has achieved approximately 100% accuracy for TCR construction (Supplementary Fig. 7A, 7B). This finding underscores the adequacy of a 150bp read length for this task. Notably, a 100bp read length falls short for TCR construction in 5’ scTCR-seq libraries. In such cases, only J genes can be reliably identified, due to their proximity to the 5’ of sequencing reads (Fig. 3A).

Furthermore, we explored the significance of sequencing depth on the performance of the methods. Initially, we compared the sequencing depth differences between cells with and without AsTCRs in experimental scRNA-seq data. Cells with correct AsTCRs displayed deeper sequencing depth (Fig. 3B and Supplementary Fig. 8A). To estimate the TCR sequencing depth in experimental scRNA-seq data, we calculated TCR contig depths using TRUST4 as its outstanding performance (Fig. 3C). Based on this estimation, we simulated data with varying sequencing depths, ranging from 2X to 100X. The results demonstrated that most methods exhibited higher TCR assembly accuracy with increased sequencing depth. However, we observed distinct patterns for successfully assembled ratios under varying sequencing depths (Fig. 3D-G and Supplementary Fig. 8B, C). For instance, TRUST4 and TraCeR achieved optimal results at a sequencing depth of 2X, while MiXCR required deeper sequencing depths of approximately 100X (Fig. 3D-G and Supplementary Fig. 8B, C). In contrast, DeRR demonstrated robust performance in assembly results at a sequencing depth of 400X (Supplementary Fig. 7A, B). Conversely, CATT and ImRep demonstrated suboptimal performance in the simulated data. These findings underscore the significance of sequencing depth for TCR construction and provide a filtering cut-off criterion for users seeking successful TCR assembly from scRNA-seq data. Specifically, for TRUST4, scRNA-seq data with a total read count exceeding 100,000 per cell or TCR sequencing depth greater than 2 are more likely to result in successful TCR assembly.

### Performance Analysis of Different Methods Using scTCR-seq and Bulk TCR-seq Data

The performance of most methods was notably superior on simulated data compared to experimental data. This discrepancy can largely be attributed to the limited presence of TCR-related reads in the experimental data. Due to TCR-specific amplification, (sc)TCR-seq data typically contains adequate TCR-related reads. In addition, we also noted that these methods can handle TCR assembly from (sc)TCR-seq data. Therefore, we also assessed the TCR assembly performance on (sc)TCR-seq data (Supplementary Table 2).

For scTCR-seq data, MiXCR displayed sensitivity and accuracy close to 100%, while the results of TRUST4 and DeRR were slightly inferior (Fig. 4A-D). In the assessment of bulk TCR-seq data, TRUST4 and MiXCR assembled the most number of CDR3β and also showed the most overlaps between them (Supplementary Fig. 9A-D).

**Figure 4.**
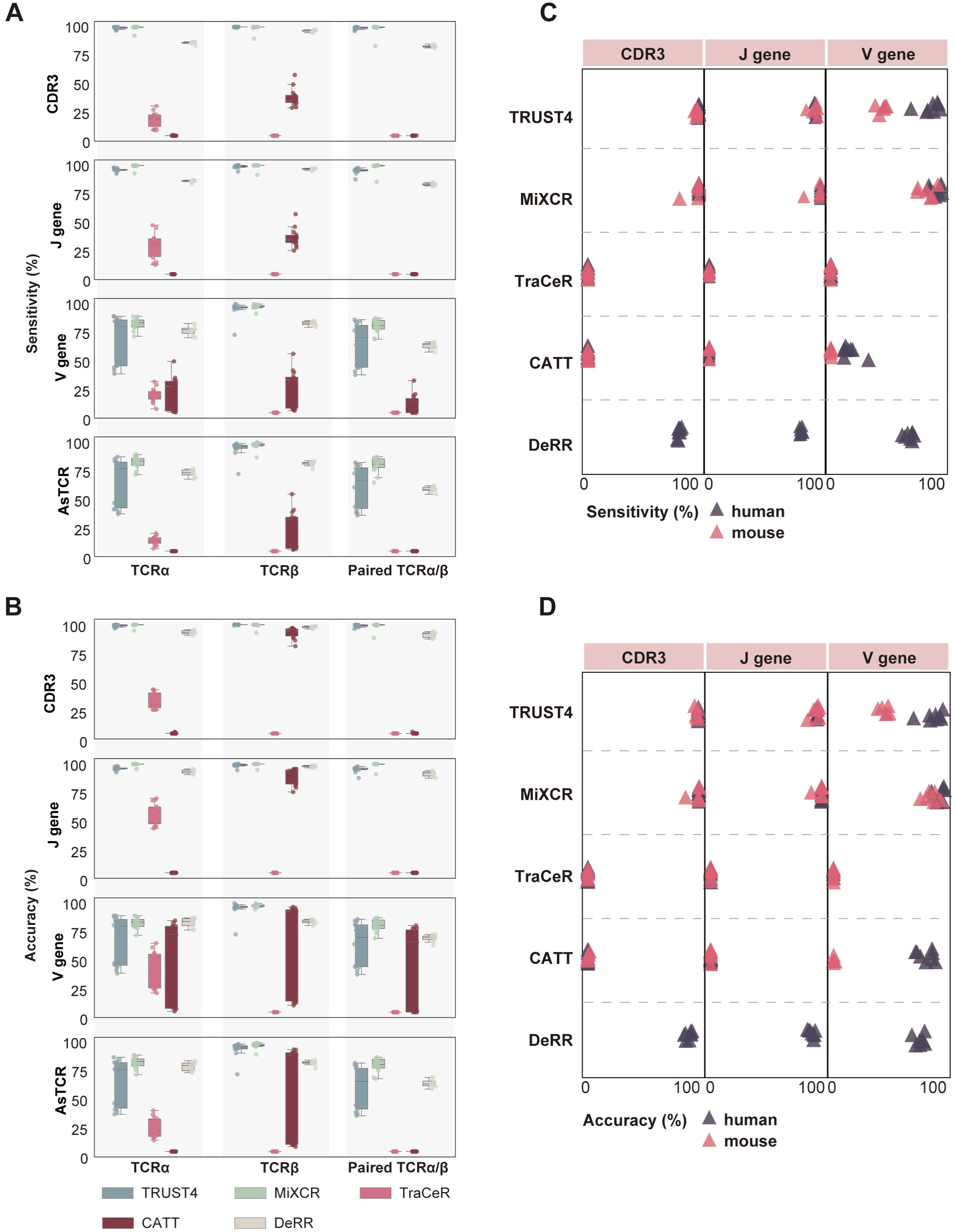
Comparative analysis of TCR construction methods on scTCR-seq data. **A.** Box plot represents the sensitivity of TCR prediction of different methods. The line in the middle represents the median; boxes represent the 25th (bottom) and 75th (top) percentiles; and whiskers represent the minimum and maximum points within 1.5 times the interquartile range. **B.** Box plot represents the accuracy of TCR prediction of different methods **C-D.** Dot plot represents the sensitivity **(C)** and accuracy **(D)** of different methods for mouse and human data.

In summary, TRUST4 and MiXCR demonstrated commendable performance in TCR assembly for both scTCR-seq and bulk TCR-seq data. They proved to be suitable not only for scRNA-seq data with limited TCR-related reads but also for TCR-seq data with abundant TCR-related reads.

### Performance Analysis of Different Methods Using Pseudo Bulk RNA-seq with Varying Cell Numbers

In light of the substantial amount of available bulk RNA-seq data and the crucial role of bulk T-cell repertoire analysis in diverse diseases [38–40], it is therefore essential to evaluate the performance of these TCR construction methods on bulk RNA-seq data. Consequently, we generated pseudo-bulk RNA-seq data derived from experimental scRNA-seq data to assess the performance of these methods across varying cell numbers, including 100, 500, and 1000 cells. TRUST4 exhibited exceptional sensitivity in both scRNA-seq and bulk RNA-seq data. Conversely, MiXCR demonstrated heightened sensitivity and accuracy specifically for bulk RNA-seq data in contrast to scRNA-seq. CATT displayed notable sensitivity, particularly in the TCRβ. Furthermore, DeRR showcased highest accuracy in the identification for both CDR3α and CDR3β, while its sensitivity remains comparatively modest concerning other methods (Fig. 5). Generally, our findings revealed that most methods displayed increased accuracy and sensitivity as the cell number increased (Fig. 5). Moreover, similar to the observations in single-cell sequencing data, β chains tend to exhibit higher accuracy than α chains in TCR assembled from bulk RNA-seq data (Fig. 5).

**Figure 5.**
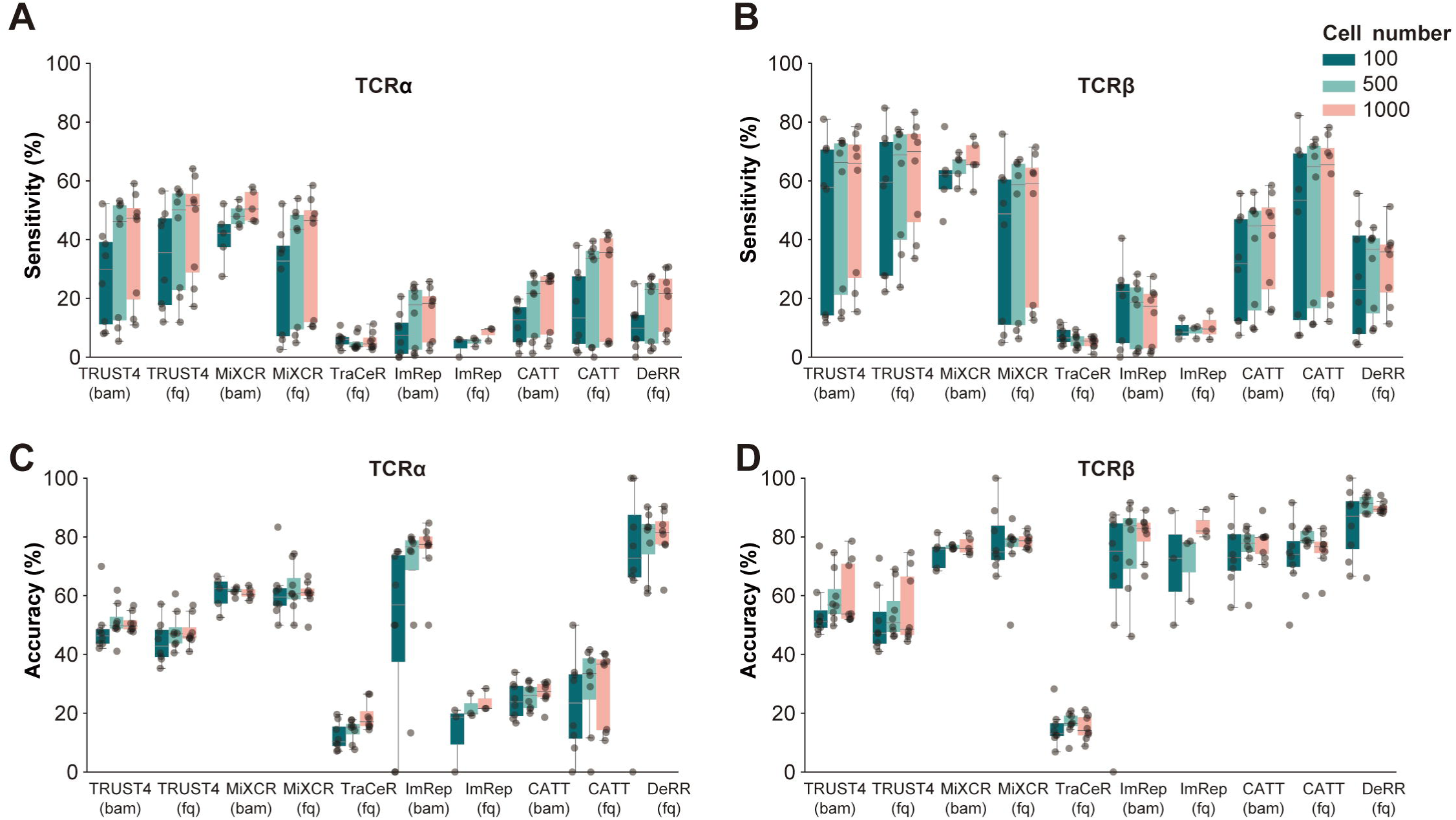
Performance assessment of different methods on pseudo bulk RNA-seq with varying cell numbers A-D. Box plots represent the sensitivity **(A, B)** and accuracy **(C, D)** of TCRα and TCR3β assembled by different methods at varying cell numbers. The line in the middle represents the median; boxes represent the 25th (bottom) and 75th (top) percentiles; and whiskers represent the minimum and maximum points within 1.5 times the interquartile range. fq, FASTQ format; bam, Binary Alignment Map format.

TCR abundance may be a crucial factor influencing the correct assembly of TCRs. Therefore, we evaluated the sensitivity and accuracy of pseudo bulk RNA-seq with varying TCR abundance (Supplementary Fig. 10). We categorized cells in each sample into three equal-sized groups (high, medium, low) based on their TCR abundance, and compared the sensitivity and accuracy among different groups. The results indicate that higher TCR abundance is associated with increased performance in most cases. Additionally, TRUST4 and MiXCR consistently performed relatively well across different TCR abundance levels compared to other methods.

### Overall Scoring and Ranking of Methods

In the evaluation of the diverse methods, we adhered to the established guidelines for benchmarking computational biology-related methods [41]. To comprehensively assess their performance, we considered six key aspects, namely accuracy, sensitivity, adaptability, usability, time consumption, and memory usage. A quantitative composite score was then computed for each evaluated method (see Methods section). TRUST4 displayed the highest overall accuracy, while DeRR exhibited the highest average sensitivity across all tested experimental scRNA-seq data (Fig. 6A, B). Regarding memory and time consumption, TRUST4 and DeRR exhibited more favorable performance in memory, with TRUST4 displaying the best performance in time (Fig. 6C, D). Additionally, the memory and time consumption of these software tools fall within the acceptable limits for most computational resources. By assigning appropriate weights (0.2 for accuracy and sensitivity, and 0.1 for other aspects), we calculated an overall score for each method, subsequently ranking them accordingly (see Methods). TRUST4 achieved the highest score (6.97), followed by MiXCR (4.79), DeRR (4.47), ImRep (2.55), TraCeR (2.39), CATT (2.29) and BASIC (1.98) in descending order (Fig. 6E-G).

**Figure 6.**
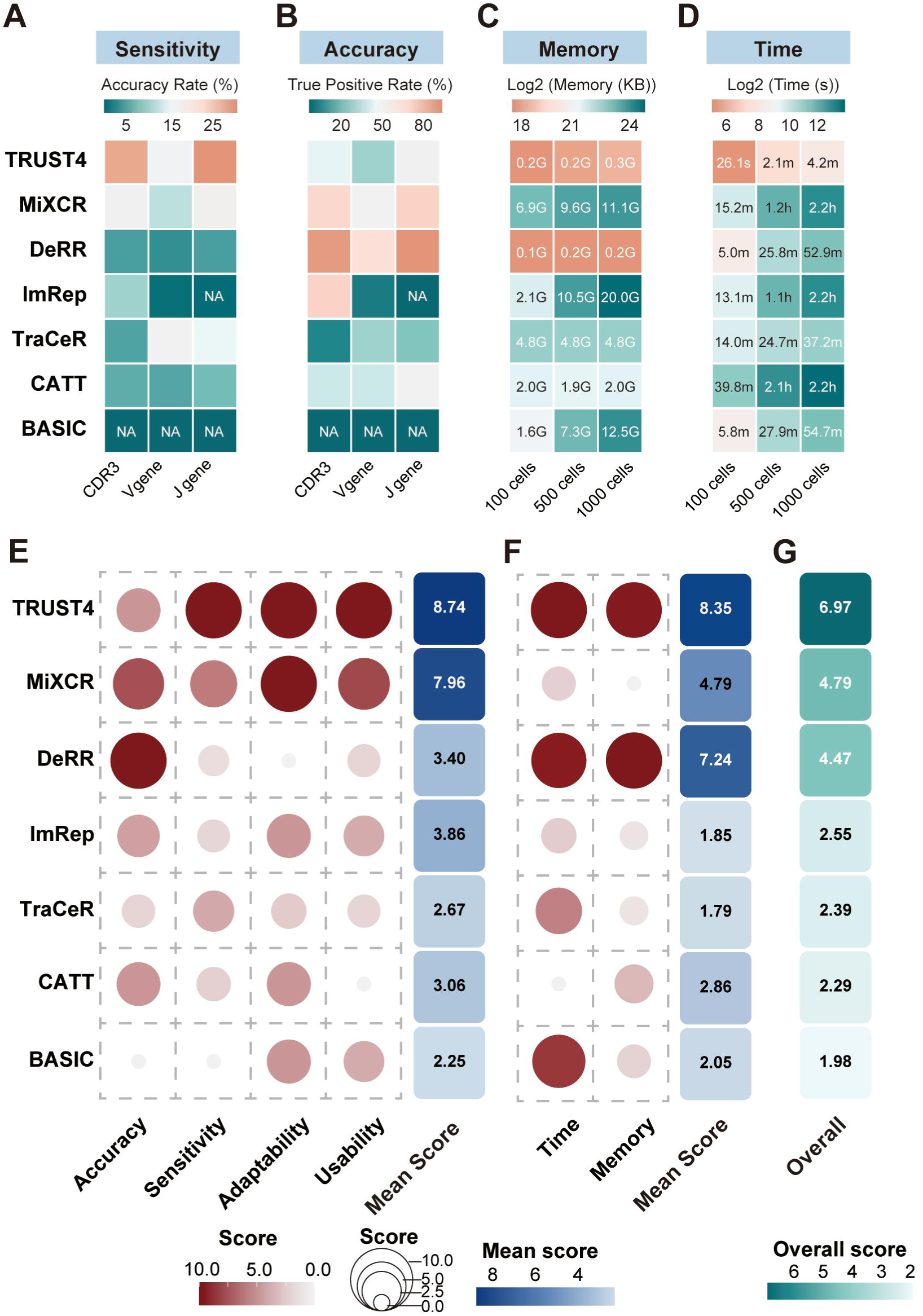
Comprehensive scoring and ranking for evaluating methods performance A-D. Heatmap represents the comparison of the sensitivity **(A)**, accuracy **(B)**, memory usage **(C)**, and time usage **(D)** of TCR reconstruction performed by different methods. NA, not available; G, gigabytes; h, hours; m, minutes; s, seconds. **E.** Dot plot represents the comprehensive evaluation of each method’s performance, measured using a min-max scale, where a higher score indicates better performance across accuracy, sensitivity, adaptability, and usability. The heatmap represents the mean scores. **F.** Dot plot represents the comprehensive assessment of each method’s computational consumption, also using a min-max scale, with higher scores signifying better performance in terms of time and memory efficiency. The heatmap represents the mean scores. **G.** Heatmap represents the overall score and ranking of each method and was presented in descending order. A weight of 0.2 was assigned to sensitivity and accuracy, while a weight of 0.1 was assigned to the other factors considered.

## DISCUSSION

In this study, we conducted an extensive and rigorous benchmark analysis to evaluate the performance of seven TCR construction methods using (sc)RNA-seq datasets. We opted to exclude VDJPuzzle [26] from our evaluation since it only supports paired-end data types and its excessive time consumption. Leveraging experimental single-cell immune profiling datasets, along with pseudo-bulk RNA-seq, and simulated scTCR-seq datasets, we systematically assessed the accuracy and sensitivity of the TCR construction process, particularly in recovering CDR3 and VJ genes for both TCRα/β chains. This comprehensive assessment encompassed diverse factors such as sequencing depths, read lengths, library construction strategies, and input data types. Additionally, we have also assessed the computational performance of each method. Overall, our study generated a guideline that can aid researchers in selecting TCR construction methods for their specific research needs.

Previous studies have benchmarked the performance of various methods for BCR construction from SMART-seq data [40]. However, the generation of BCR involves somatic hypermutations and isotype switching, which are not observed in TCR generation [4,41]. Thus, the conclusions of BCR construction methods may not be directly applied to TCR. Therefore, it is essential to independently benchmark the methods for TCR construction, as highlighted in the BCR benchmark study [42]. In this study, we also utilized the datasets generated by the more popular 10X strategy and developed a scTCR simulator, YASIM-scTCR which enables us to precisely analyze the accuracy and sensitivity of different methods.

In the context of 10X scRNA-seq data, our analysis revealed that TRUST4 and MiXCR exhibited the highest sensitivity, followed by CATT and TraCeR, while DeRR and MiXCR demonstrated superior accuracy (Fig. 2A, C). However, when considering bulk RNA-seq data, which generally offers deeper sequencing depths compared to scRNA-seq data, a broader range of methods including TRUST4, MiXCR, and CATT demonstrated favorable performance (Fig. 5A-D). In a previous study, TRUST4 exhibited suboptimal performance for constructing BCR from scRNA-seq datasets [29]. This discrepancy could be attributed to the intrinsic differences observed in the TCR or BCR recombination process or the usage of 10X scRNA-seq data which may be better suited for TRUST4 [29]. Overall, the choice of TCR construction methods should be made judiciously, considering the specific characteristics of the data and the research objectives at hand.

In this study, we placed particular emphasis on the influence of candidate reads, a critical factor affecting TCR construction. To facilitate comprehensive analyses, we developed a novel simulator called YASIM-scTCR, enabling the generation of scTCR-seq data with TCR- and non-TCR-derived reads, accommodating user-defined parameters such as sequencing depths and read lengths. For BCR data analysis, there is a dedicated simulator called AIRRSHIP, which could potentially provide more comprehensive and detailed insights for BCR benchmark studies [43]. Through simulations with YASIM-scTCR, we observed a clear association between candidate read performance and TCR construction accuracy, where methods displaying superior candidate read identification achieved higher accuracy, while those with suboptimal candidate read identification exhibited lower accuracy (Supplementary Fig. 5A-B). This underscores the pivotal role of candidate reads in the TCR construction process. However, it is important to note that the definition of candidate reads varies across different methods, lacking a unified standard, which may introduce bias in our interpretation. Nevertheless, we believe that accurately identifying TCR-related reads is fundamental for achieving optimal results in subsequent assembly processes.

We have summarized the algorithm employed by each method (Supplementary Table 5). It is evident that those who implement TCR-specific alignment algorithms (e.g., TRUST4, and MiXCR) yield better performance than those who rely on general-purpose aligners (e.g., CATT) or more naïve algorithms (e.g., ImRep). In addition, assemblers with more advanced clustering algorithms were more likely to yield better results (e.g., TRUST4, and MiXCR), whereas those relying on V/J gene calling trend behave poorly (e.g., TraCeR, CATT, and ImRep). Even though not benchmarked in this study, we also noticed that certain methods use expectation-maximization (EM) algorithms in quantification (e.g., TRUST4, and TraCeR), which should theoretically improve clonal expansion quantification accuracy for bulk TCR-seq data.

Using experimental datasets, our analysis suggested that the performance of most methods was not significantly influenced by human and mouse. However, a notable discrepancy in performance emerged between single-end and paired-end sequencing data. Thus, we recommend using the paired-end mode (5’ library construction strategy) for more accurate TCR construction (Supplementary Fig. 6A-D). Regarding read length, our simulation analysis demonstrated that it does not exert a critical influence on TCR construction. In contrast, sequencing depth emerged as a crucial determinant of accuracy. This emphasizes once again the critical importance of sequencing depth in NGS experiments [44,45].

Our investigation has shed light on certain limitations of TCR construction from scRNA-seq data. Notably, we observed that the performance of methods on simulated data outperformed that on experimental scRNA-seq data, with TRUST4 being a prominent example. In the simulated data, TRUST4 achieved remarkable accuracy in TCRβ chain assembly (Supplementary Fig. 5B), whereas its accuracy was dramatically reduced on experimental scRNA-seq data (Fig. 2A). This discrepancy may be attributed to the fact that the simulated data encompassed abundant TCR-related reads, which is not true for experimental scRNA-seq data due to factors like limited sequencing depth or suboptimal library construction strategies. To overcome these limitations, utilizing alternative sequencing methods, such as adopting more in-depth sequencing or employing paired-end mode (5’ library construction strategy), may address these challenges and enhance the accuracy of TCR construction from scRNA-seq data.

When assessing the performance of various methods using experimental scRNA-seq data, we employed the output of CellRanger from scTCR-seq as our ground truth. scTCR-seq involves specific amplification of TCR sequences, making it a reliable source for ground truth data. While our results indicated that the performance of MiXCR, TRUST4, and DeRR are largely consistent with CellRanger on experimental scTCR-seq data, it is important to note that CellRanger itself is a method for single-cell TCR construction, which may introduce potential bias. In addition, simulated data with YASIM-scTCR provide an accurate ground truth without the potential bias introduced by CellRanger.

In summary, we conducted a comprehensive evaluation of seven distinct methods across six aspects (Fig. 6A-G). Among these methods, TRUST4, MiXCR, and DeRR emerged as the top performers, with TRUST4 demonstrating superior sensitivity, and DeRR excelling in accuracy (Fig. 6A-G). Furthermore, both TRUST4 and DeRR exhibited commendable efficiency concerning time and memory consumption. The outcome of our evaluation thus furnishes users with valuable recommendations for selecting appropriate methods tailored to their specific needs, and it also guides developers with valuable insights to enhance and optimize the performance of their methods.

## Materials and Methods

### Datasets and preprocessing

14 paired 10X scRNA-seq and scTCR-seq datasets comprising both paired-end and single-end reads from human and mouse were collected in this study (Supplementary Table 2). The scTCR-seq data was analyzed using CellRanger (version 6.1.2) by “cellranger vdj” command as the ground truth for each cell, while the scRNA-seq data was analyzed using CellRanger by “cellranger count” command and alignment files in BAM format was obtained. Given that only TRUST4 and MiXCR support the 10X format, while other software do not, we chose to split the input data into individual cells to standardize the input. This approach ensured compatibility across all software for evaluation. To further construct TCR from scRNA-seq data, we split the data in both FASTQ and BAM format into individual cells using a Python script, which is available on GitHub. Furthermore, we collected a total of 14 bulk TCR-seq datasets from both human and mouse and 50 SMART-seq datasets from mouse, with details provided in Supplementary Table 2. All datasets are publicly available for download from the National Center for Biotechnology Information (NCBI).

### TCR Construction Using Various Methods

All software was installed on Ubuntu 20.04.4, following the installation instructions and documentation provided by each software. We referred to the software’s user manual to determine the appropriate parameters for each run, and specific software versions and parameter settings used are detailed in Supplementary Table 3.

### The Definition of Accuracy and Sensitivity

For each experimental sample, “all cells” are cells inside the “filtered_contig_annotations” output from CellRanger, whose TCRs are both full-length and productive. To evaluate the performance of each method, cells without assembly results are considered “no-result cells,” and the rest are considered “result cells”. Among the result cells, if at least one of the TCRs is correct, the cell is considered to be correctly assembled (named “true cells”). For each sample, sensitivity is defined as the percentage of true cells out of all cells in the sample, while accuracy is defined as the percentage of true cells out of the result cells. Evaluation of the V/J gene is on the subgroup level with CDR3 sequences on the amino acid level. The AsTCRs are defined as those having both the V/J genes and CDR3 sequences correct.

### The Definition of Candidate Reads

In Supplemental Fig. 5, we used simulated data to explore the results for candidate reads analysis among the 4 methods: DeRR, ImRep, TraCeR, and TRUST4. For DeRR, we modified its “DeRR.py” file by removing the line “os.system(f“rm -f {sam_file} &”)” at line 217. This change allowed it to keep TRJ.sam and TRV.sam files, which were considered candidate reads in this study. For ImRep, we defined candidate reads as the reads mentioned in the files starting with “partial_cdr3” in its output. Regarding TraCeR, we considered candidate reads as the reads involved in TCR_A.fastq and TCR_B.fastq, located in the “aligned_reads” folder generated by the method. As for TRUST4, candidate reads were defined as the reads inside “toassemble.fq” file.

### The Definition of TCR Contig Depths from Experimental scRNA-seq

The calculation process for TCR contig depths is as follows: 1) Identify cells with successful TCR assembly from TRUST4. 2) Determine the TCR contig length. 3) Calculate the candidate reads from TRUST4 output in cells where TCR has been successfully assembled. 4) Compute the total number of nucleotides in candidate reads and divide it by the TCR contig length.

### Simulation of scTCR-seq and scRNA-seq Data

We employed YASIM-scTCR (version 1.0) to simulate 10X Genomics Single Cell Immune Profiling 5’ scTCR-Seq with TCR- and non-TCR-derived. YASIM-scTCR is a Python-based tool capable of generating realistic TCR recombination events by leveraging data from huARdb [14,35]. The YASIM-scTCR package is available on PYPI (https://pypi.org/project/yasim-sctcr/1.0.0/), and its source code can be accessed on GitHub (https://github.com/WanluLiuLab/yasim-sctcr).

To perform the simulations, we utilized specific reference data sources for the genome and gene annotations. The soft-masked GRCh38.p12 genome sequence was obtained in FASTA format from Ensembl (release 97) and served as the reference genome for our simulations. The corresponding cDNA and peptide sequences in FASTA format of this release were also downloaded. Genes defined on chromosomes 7 and 14 with TCR-related biotypes were extracted using “seqkit grep” (version 2.5.1) [46]. The YASIM-scTCR “generate_tcr_cache” module will then prepare the references by aligning selected cDNAs and peptides using Smith-Waterman alignment [47], which served as the basis for generating realistic TCR recombination events during the simulations.

The process of simulating TCR rearrangements is conducted using the “rearrange_tcr” module of YASIM-scTCR. Before the generation of each TCR, whether the TCR sequence would be producible is based on a fixed probability. Firstly, TRAV, TRAJ, TRBV, and TRBJ genes are selected from the V/J gene library using the V/J usage bias pattern from huARdb (Supplementary Fig. 4). Then, the V/J gene clipping on the TRA and TRB chains is performed based on statistics from huARdb (Supplementary Fig. 3A, B). Subsequently, CDR3 sequences are generated according to the statistics from huARdb (Supplementary Fig. 3C). Furthermore, V/J genes are subjected to additional clipping criteria to ensure that the amino acid of the V gene starts with C (cysteine) and the J gene ends with F (phenylalanine, for most genes) or W (tryptophan, for TRAJ33, TRAJ38, TRAJ55 only). In cases where the generation fails, the simulator retries until successful, ensuring the production of reliable and realistic TCR sequences. Approximately 50bp of the corresponding C gene segment is then added, mimicking the behavior of C-specific primers in scTCR-seq. The generated ground-truth TCR contigs are exported in FASTA format, preserving the ground-truth V/J gene names and CDR3 sequences.

Even though not considered in our simulated data, YASIM-scTCR also supports simulation for TCR clone expansion level when generating a large amount of scTCR-seq data. After the introduction of TCR clonal expansion (Zipf’s distribution) and sequencing depth (uniform distribution), the mRNA contigs were reverse complemented to allow 5’ amplification using seqtk [48]. Finally, driven by YASIM-scTCR, ART [36] with “--amplicon” parameter is utilized with desired sequencing length and machine errors, providing simulated 5’ scTCR-seq data containing TCR-derived reads.

The simulation of scTCR-seq data with non-TCR-derived reads involves several steps. Firstly, the reference mRNAs are filtered for protein-coding genes (with “protein_coding” biotype) selected by Matched Annotation from NCBI and EMBL-EBI (MANE) [49]. For expression data, we used scDesign2 [36] trained by HU_0043_Blood_10X dataset from the Human Universal Single Cell Hub (HUSCH) database on HUGO Gene Nomenclature Committee (HGNC)- approved MANE-selected protein-coding genes [50,51]. YASIM-scTCR could also accept count matrices in other formats that could be parsed by AnnData [52]. The expression data were manually scaled to a mean depth of 1 for each gene per cell. All transcripts were reverse-complemented and passed to YASIM-scTCR. Additionally, each cell was barcoded to facilitate downstream analysis and identification.

### Pseudo-bulk RNA-seq for TCR Construction Analysis

We randomly selected 100, 500, and 1000 individual cells from the scRNA-seq FASTQ and BAM format data. These selected cells were combined to create pseudo-bulk RNA-seq data. BAM files were merged using SAMtools (version 1.15.1), while the FASTQ files were concatenated using the command “cat”. During the evaluation process, we recorded the time and memory consumption for each method using GNU Time.

### Scoring Principles and Standards

The evaluation scores presented in Fig. 6E are derived from six attributes, as detailed in Supplementary Table 4.

For the accuracy score, the proportion of cells correctly assembled for CDR3, V gene, and J gene in each 14 samples was calculated and averaged, with weights of 0.5, 0.3, and 0.2 respectively. To transform the accuracy score into a more interpretable range, it was scaled to a 1-10 scale using the min-max scaling method. Sensitivity scores were calculated using a similar way.

The time and memory scores for each method were calculated using (log(1/time (s)) and log(1/memory (KB))) with varying cell numbers, and scaled to 1-10. Higher scores indicate better time and memory performance. The details of the time, memory, and CPU usage are available in Supplementary Table 4.

The adaptability score was determined by assessing the number of input formats supported by each method. Various aspects, including object cell type (T/B cell), input data format (FASTQ/BAM files), and species supported (human or mouse). One point was awarded for each aspect, and the overall adaptability score was scaled to a 1-10 range.

Usability evaluation focused on two aspects: the quality and variety of the method’s output schema and the user-friendliness of the method. Four output schemas (candidate reads, assembled reads, contig sequence, annotation contig) were considered, with one point awarded for each schema supported. Additionally, the user-friendliness score, based on subjective evaluation, was assigned a maximum of 5 points. They were combined to obtain the overall usability score, which was then scaled to a 1-10 range.

The min-max scale formula is shown below:

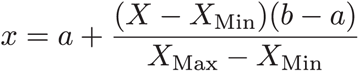

## Authors’ Contributions

**Ruonan Tian**: Conceptualization, Methodology, Software, Formal analysis, Visualization, Writing - Original Draft. **Zhejian Yu**: Conceptualization, Methodology, Software, Writing - Original Draft. **Ziwei Xue**: Data Curation, Writing - Review & Editing. **Jiaxin Wu**: Software. **Lize Wu**: Writing - Review & Editing. **Shuo Cai**: Visualization. **Bing Gao**: Data Curation. **Bing He**: Funding acquisition. **Yu Zhao**: Funding acquisition. **Jianhua Yao**: Funding acquisition. **Linrong Lu**: Funding acquisition. **Wanlu Liu**: Conceptualization, Supervision, Project administration, Funding acquisition, Writing - Review & Editing. All authors have read and approved the final manuscript.

## Data Availability

In this study, a summary of the 11 methods is provided in Supplementary Table 1. The detailed information of the public datasets in this study is available in Supplementary Table 2. All datasets can be accessed at NCBI via project accession numbers GSE114727, GSE144469, GSE160053, GSE194166, GSE223797, GSE225183, GSE74923, PRJNA393498, PRJNA412649. The command line for the 7 selected methods used in this study is presented in Supplementary Table 3. Detailed scoring information for the 7 methods across six aspects (adaptability, usability, accuracy, sensitivity, time, and memory) is presented in Supplementary Table 4. The summary of the algorithm employed by different methods in this study is available in Supplementary Table 5.

## Code availability

The source code of YASIM-scTCR can be accessed for research purposes at BioCode (https://ngdc.cncb.ac.cn/biocode/tool/BT7591) and Github (https://github.com/WanluLiuLab/yasim-sctcr) or PYPI (https://pypi.org/project/yasim-sctcr/1.0.0/). Documentation of YASIM-scTCR can be found at (https://labw.org/yasim-sctcr-docs/). The code for statistics and visualization in this paper can be accessed on GitHub via (https://github.com/WanluLiuLab/Benchmarking_TCR_Construction).

## Competing interests

Bing He, Yu Zhao, and Jianhua Yao are current employees of Tencent Technology (Shenzhen) Co., Ltd. All the other authors have declared no competing interests.

## Acknowledgments

The authors extend their gratitude to all researchers who contributed to the collection, analysis, and presentation of the datasets used in this study. Their valuable efforts and contributions were essential in making this research possible. We extend our appreciation to Dr. Junxin Lin for helping with the language editing. We thank all members of LabW for their helpful discussions and valuable suggestions. We would also like to thank the technical support provided by the Core Facilities, especially the ZJE server of ZJU-UoE Institute.

## Funding

This work is supported by the National Natural Science Foundation of China (grants 32370935, 31930038, 32100718, U21A20199, and 32170551 to W.L. and L.L.), the Fundamental Research Funds for the Central Universities 226-2022-00134 (to W.L.), the Tencent AI Lab Rhino Bird Research Funding (grant RBFR2022015 and RBFR2023009 to W.L.), the Pre-research Projects of Innovation Center of Yangtze River Delta, Zhejiang University (to L.W., W.L., L.L.), and the Innovative Research Team of High-level Local Universities in Shanghai (to L.L.).

## Declaration of AI and AI-Assisted Technologies in the Writing Process

We used ChatGPT-3.5 to assist with language refinement of the manuscript.

## Figure Legends

**Supplementary Figure 1.**
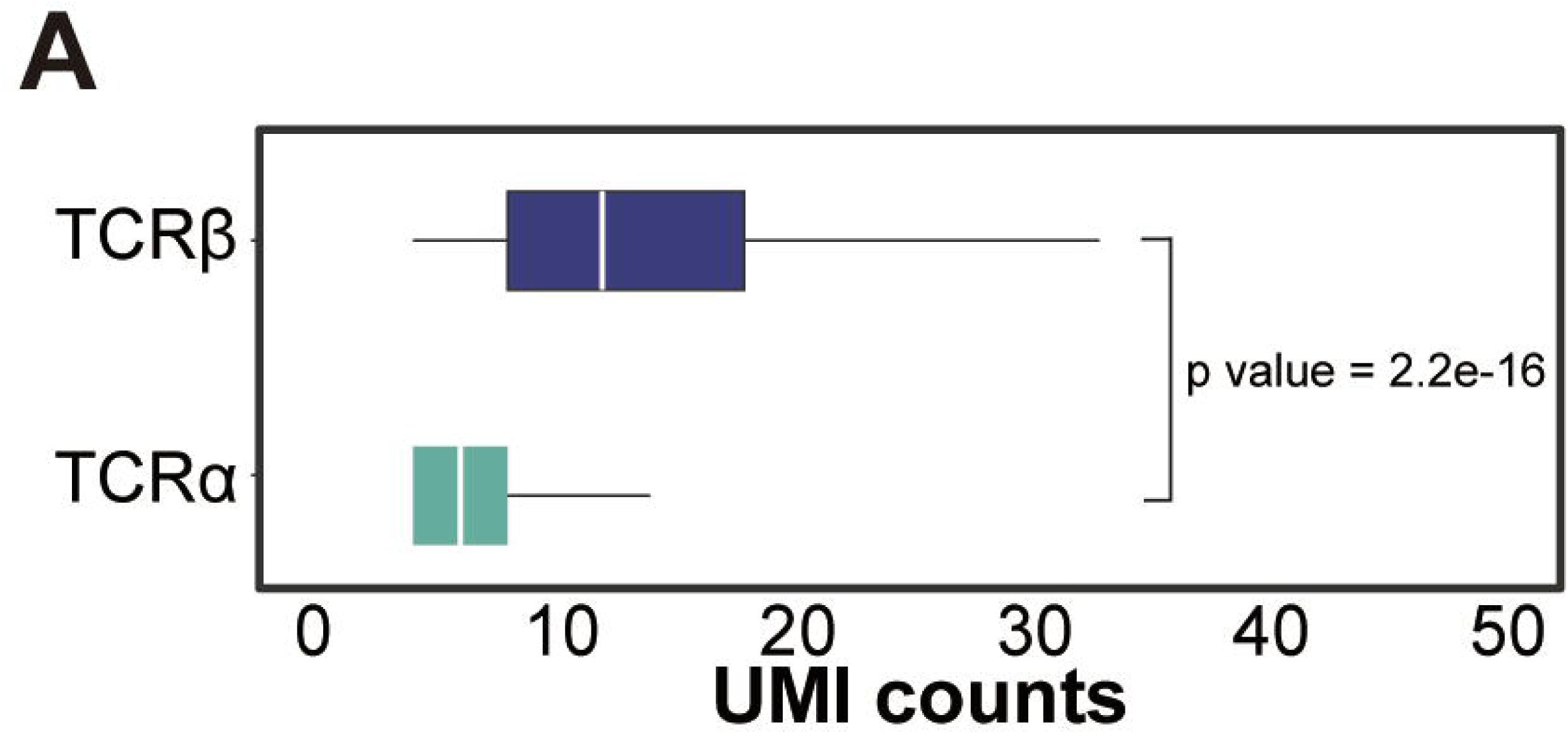
Basic statistics related to TCR. **A.** Boxplot represents the UMI counts of human CDR3α and CDR3β sourced from huARdb, accompanied by a paired Wilcoxon signed-rank test yielding a p-value of 2.2e-16.

**Supplementary Figure 2.**
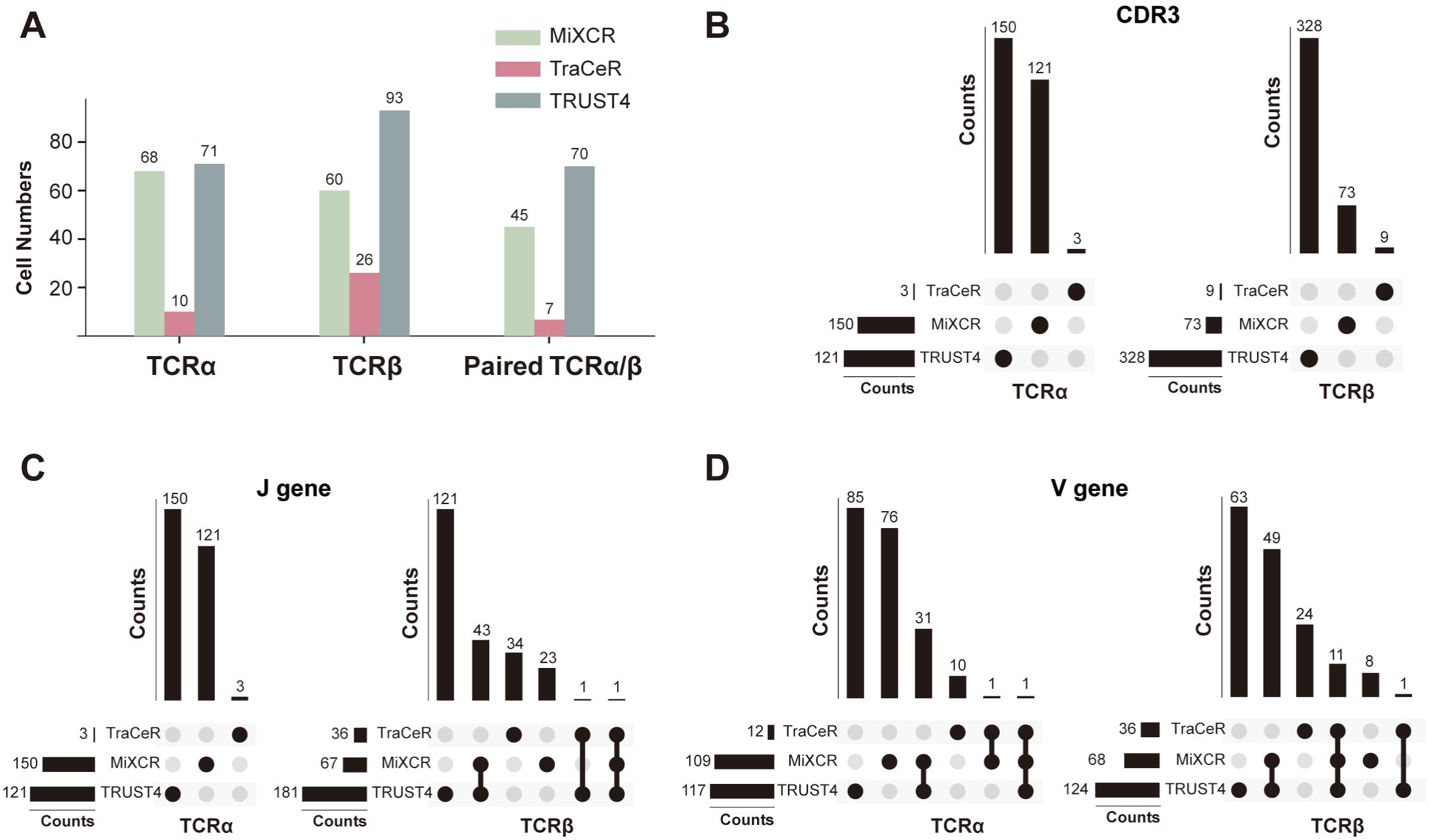
Comparative analysis of TCR construction methods on SMART-seq data. **A.** Histogram represents the number of cells for each software tool successfully reconstructed the AsTCR. **B.** Upset plots represent the overlap of unique CDR3α (left) and CDR3β (right) sequences assembled by each method. **C.** Upset plots represent the overlap of unique J gene α (left) and J gene β (right) sequences assembled by each method. **D.** Upset plots represent the overlap of unique V gene α (left) and V gene β (right) sequences assembled by each method.

**Supplementary Figure 3.**
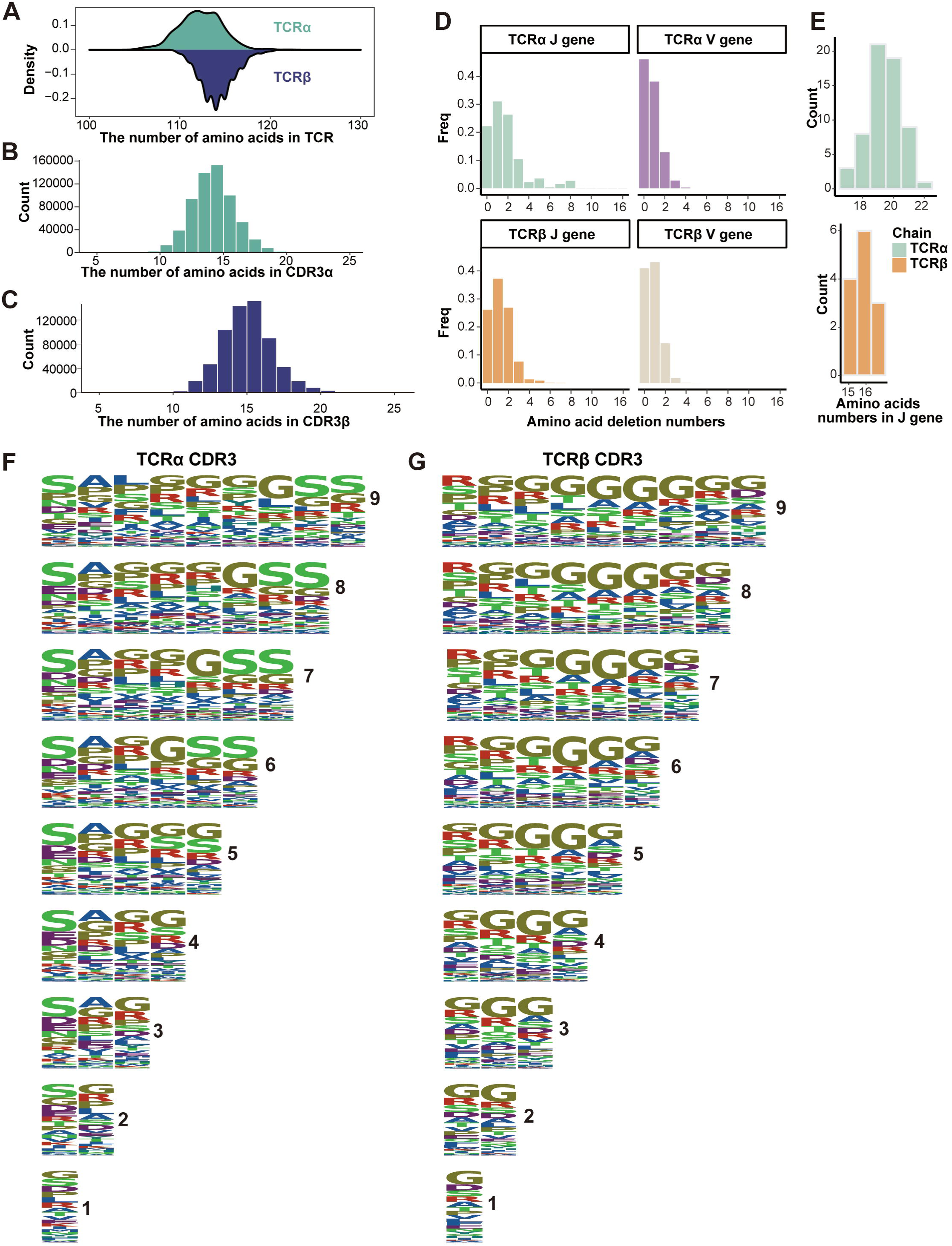
TCR CDR3 insertion and V/J gene deletion amino acid statistics. **A.** Density plot represents the amino acid length distribution of assembled human TCR sourced from huARdb. **B-C.** Histogram represents the amino acid length distribution of human CDR3α **(B)** and CDR3β **(C)** sourced from huARdb. **D.** Histogram represents the distribution of deleted amino acids number in the junction with CDR3 of V/J genes. **E.** Histogram represents the distribution of amino acid lengths of J gene. **F-G.** Logo plot of amino acid preference statistics for CDR3 of TCRα and TCRβ with different amino acid lengths. Each letter in the logo plot corresponds to a specific amino acid, represented by its single-letter code and the height of the letters in the logo plot indicates the relative frequency of each amino acid at that position.

**Supplementary Figure 4.**
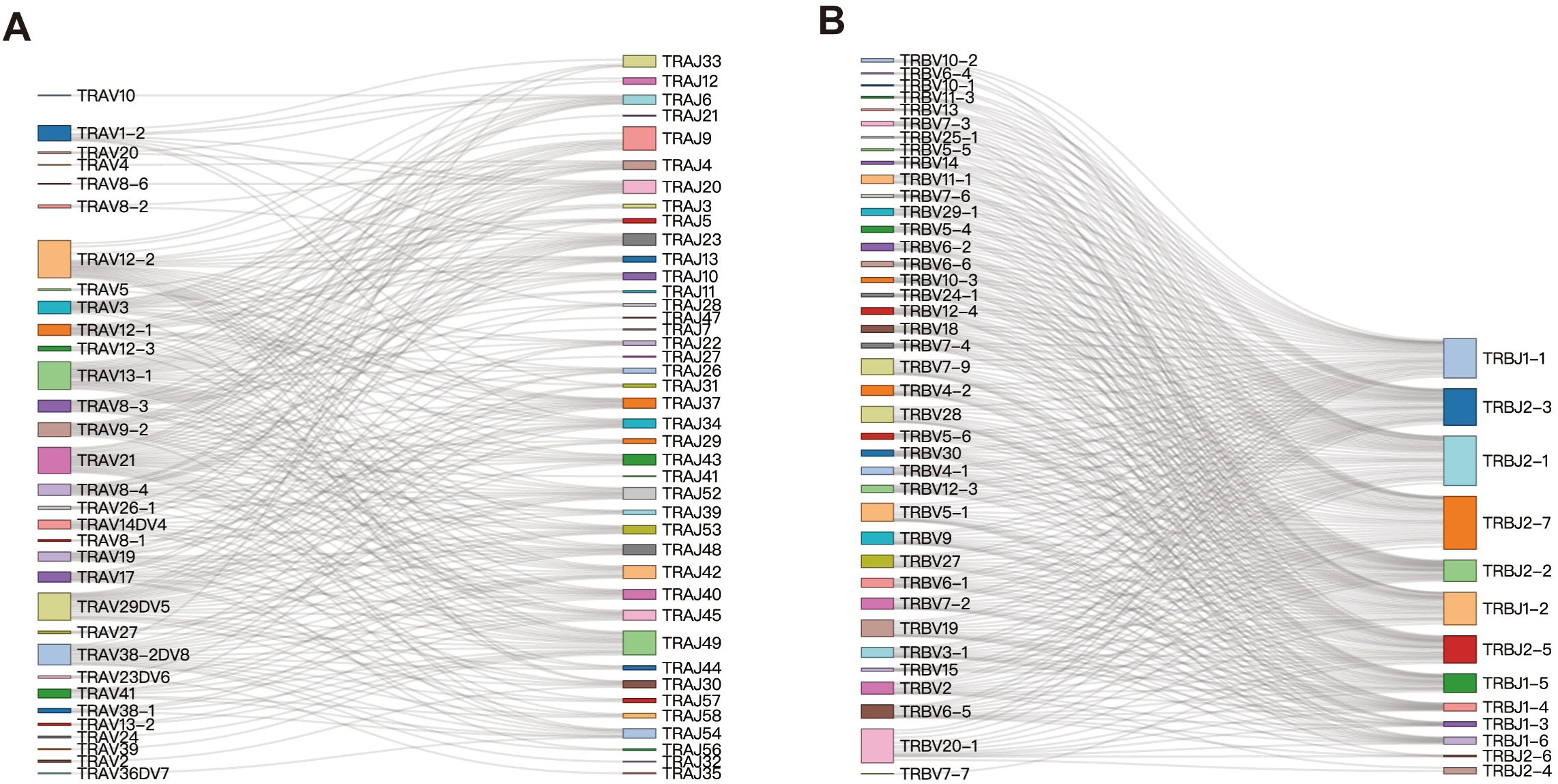
The usage of V/J gene in experimental data. A-B. Sankey plot represents the usage of V/J gene in experimental data. Each node represents each gene. Only V/J genes with a frequency greater than 0.05 were retained. Left **(A)**, TRAV gene; Right **(A)**, TRAV gene; Left **(B)**, TRBV gene; Right **(B)**, TRBJ gene.

**Supplementary Figure 5.**
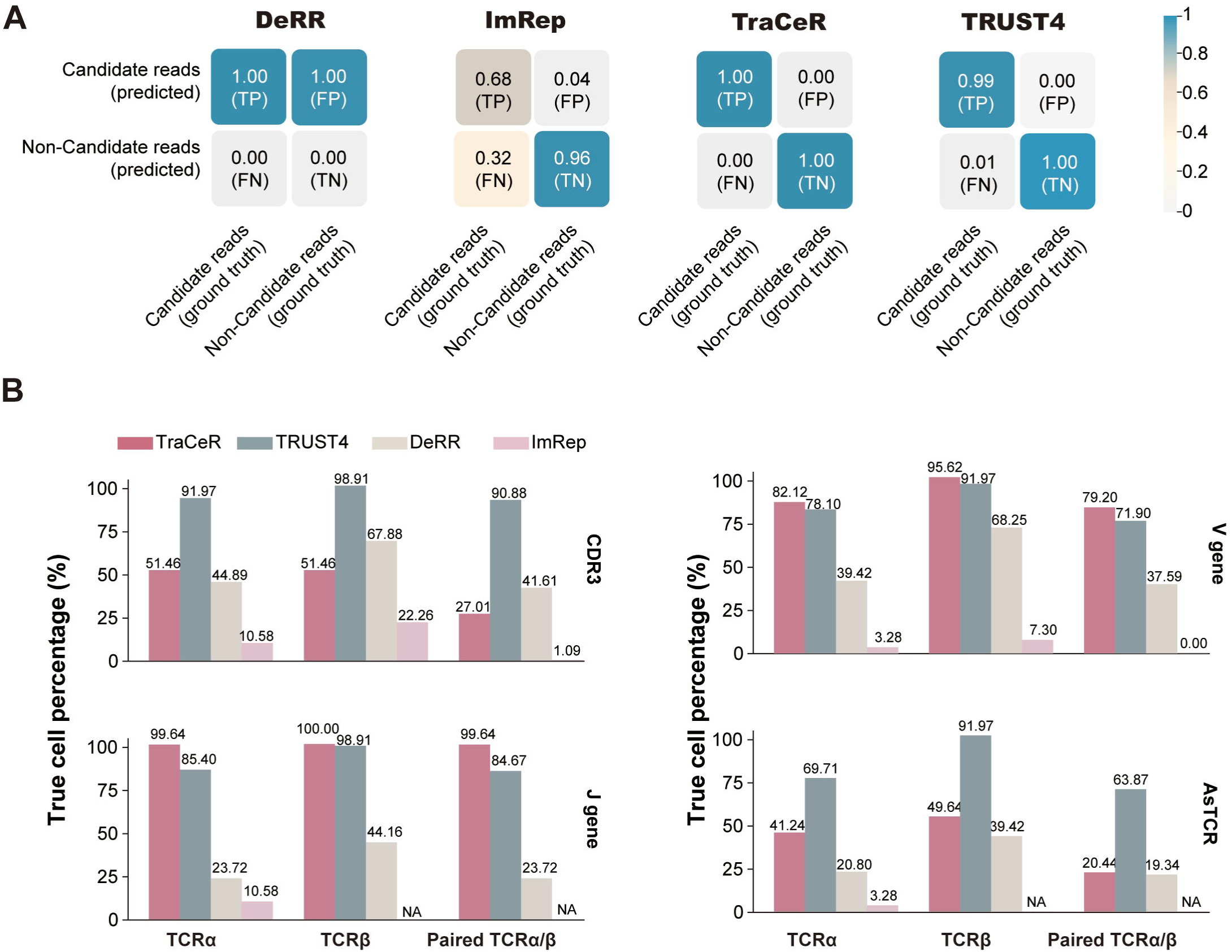
The statistics of candidate reads obtained from simulated scTCR-seq datasets. **A.** Confusion matrix for finding candidate reads in simulated scTCR-seq and scRNA-seq datasets (reads length: 250 base pair, reads depth: 400). TP, true positive; FP, false positive; FN, false negative; TN, true negative. **B.** Bar plot of four tools in terms of the accuracy of simulated datasets (reads length: 250 base pair, reads depth: 400). AsTCR, assembled TCR.

**Supplementary Figure 6.**
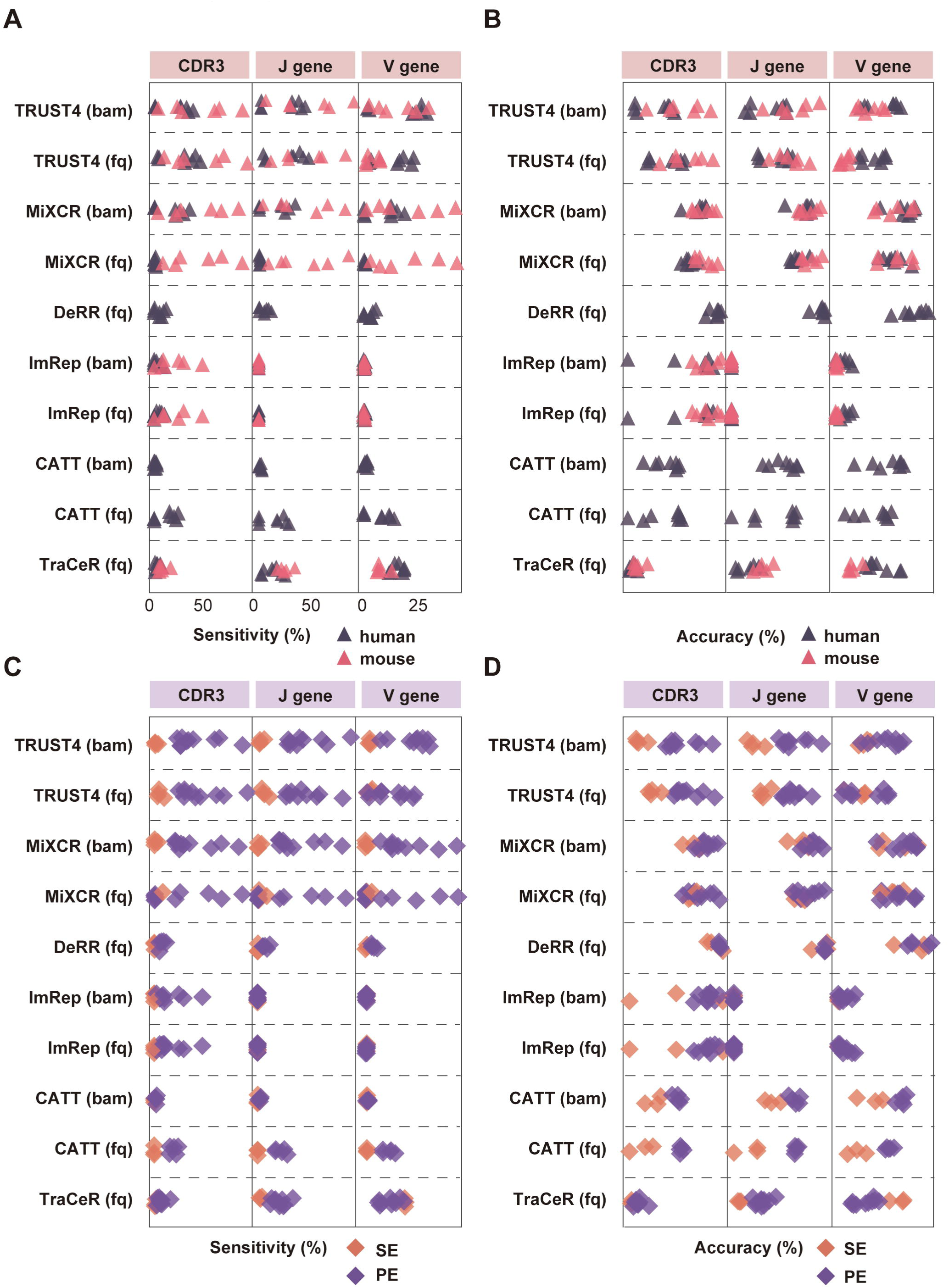
Performance comparison of methods on multiple data types A-B. Dot plot represents the sensitivity **(A)** and accuracy **(B)** of different methods for mouse and human data. **C-D.** Dot plot represents the sensitivity **(C)** and accuracy **(D)** of different methods for paired-end (PE) and single-end (SE) data.

**Supplementary Figure 7.**
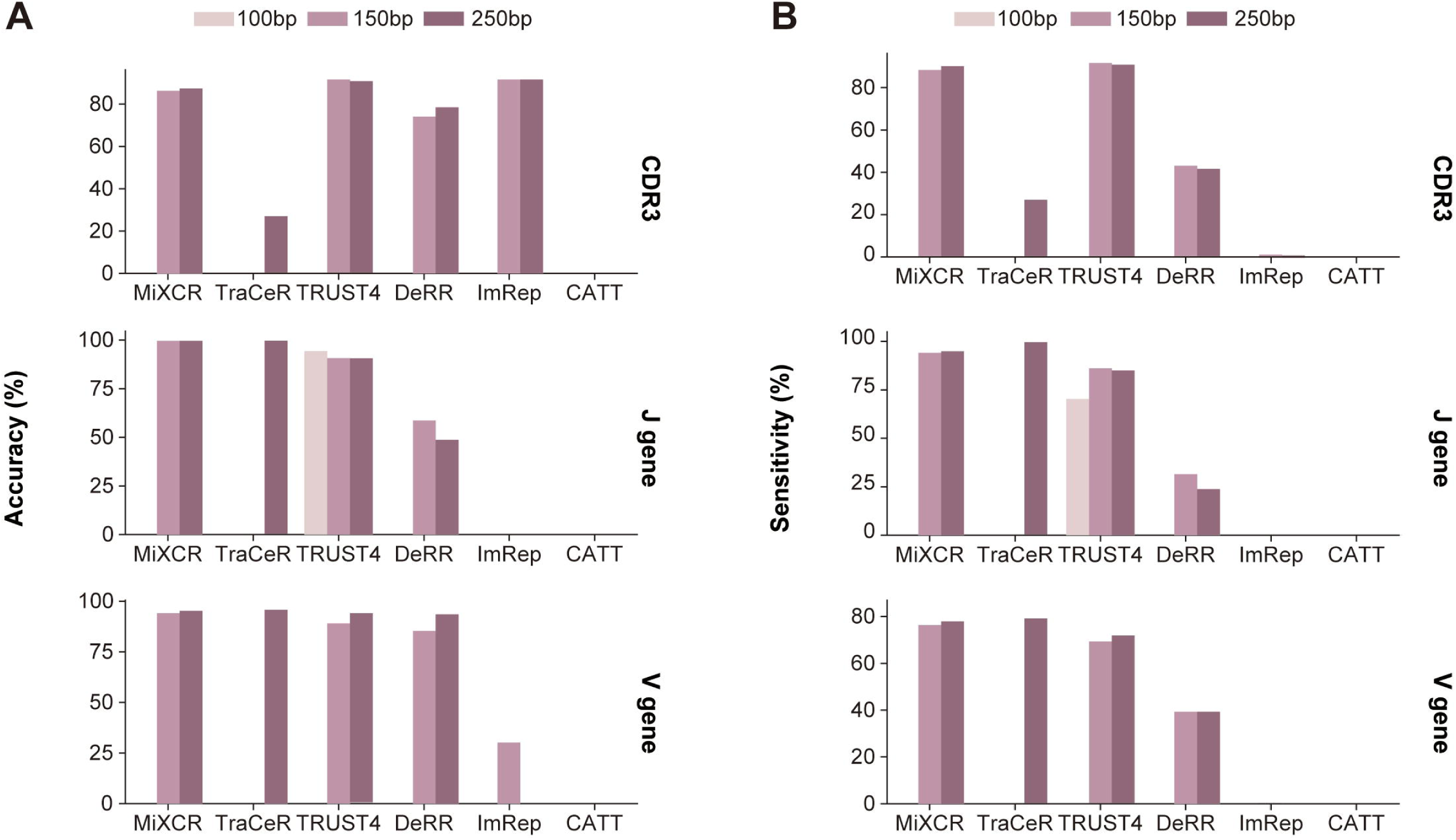
Performance comparison on simulated data with different read lengths. A-B. Bar plot represents the accuracy **(A)** and sensitivity **(B)** of TCR assembled by different software under varying read lengths.

**Supplementary Figure 8.**
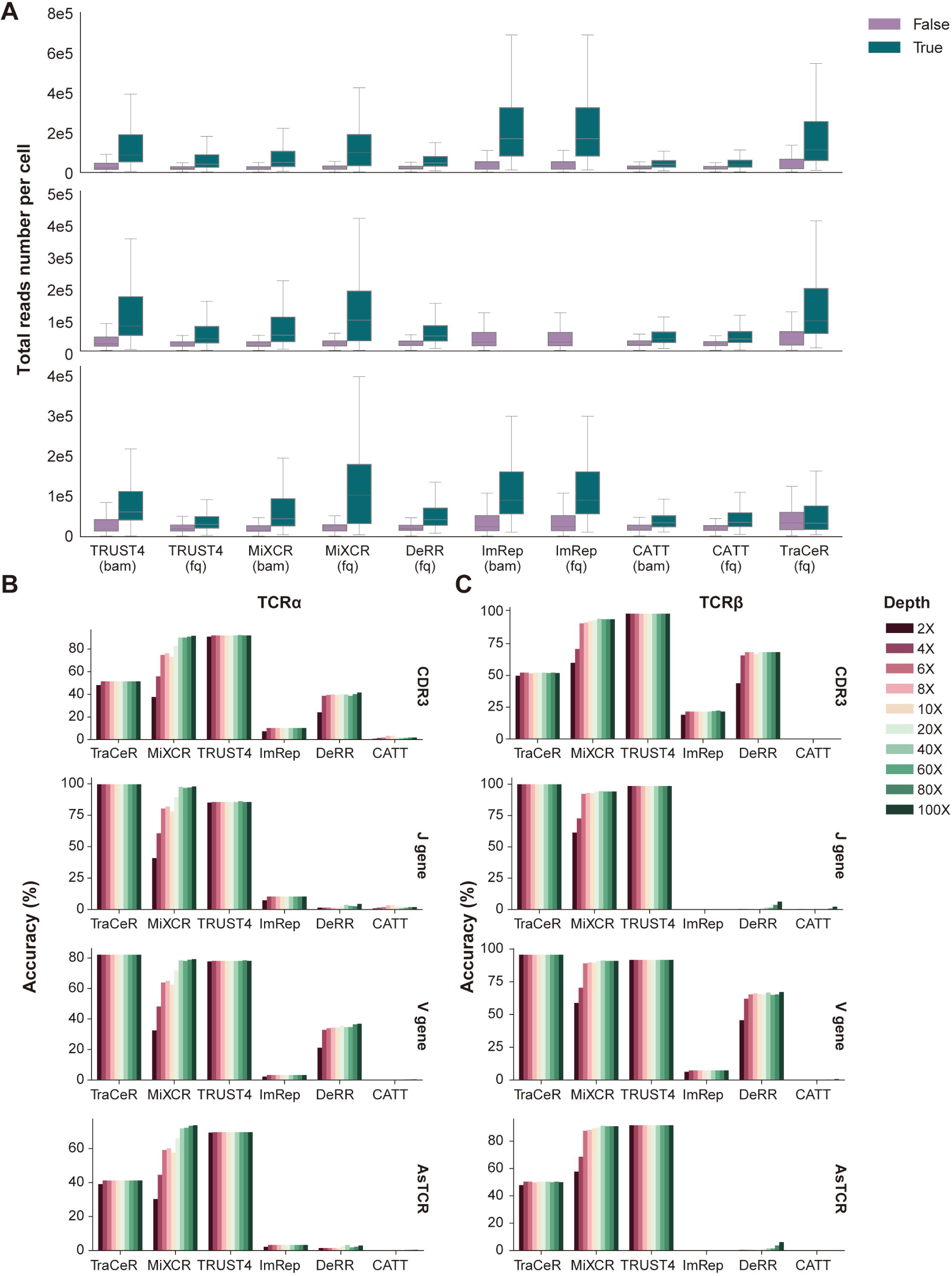
Performance comparison on simulated data with different sequencing depths. **A.** Box plot represents the total read number of CDR3, J gene and V gene in cells with successfully (True) and unsuccessfully (False) assembled TCR on the collected real scRNA-seq datasets. **B-C.** Bar plots represent the software accuracy in assembling TCRα **(B)** and TCRβ **(C)** in simulated data with different sequencing depth. AsTCR, assembled TCR; fq, FASTQ format; bam, Binary Alignment Map format.

**Supplementary Figure 9.**
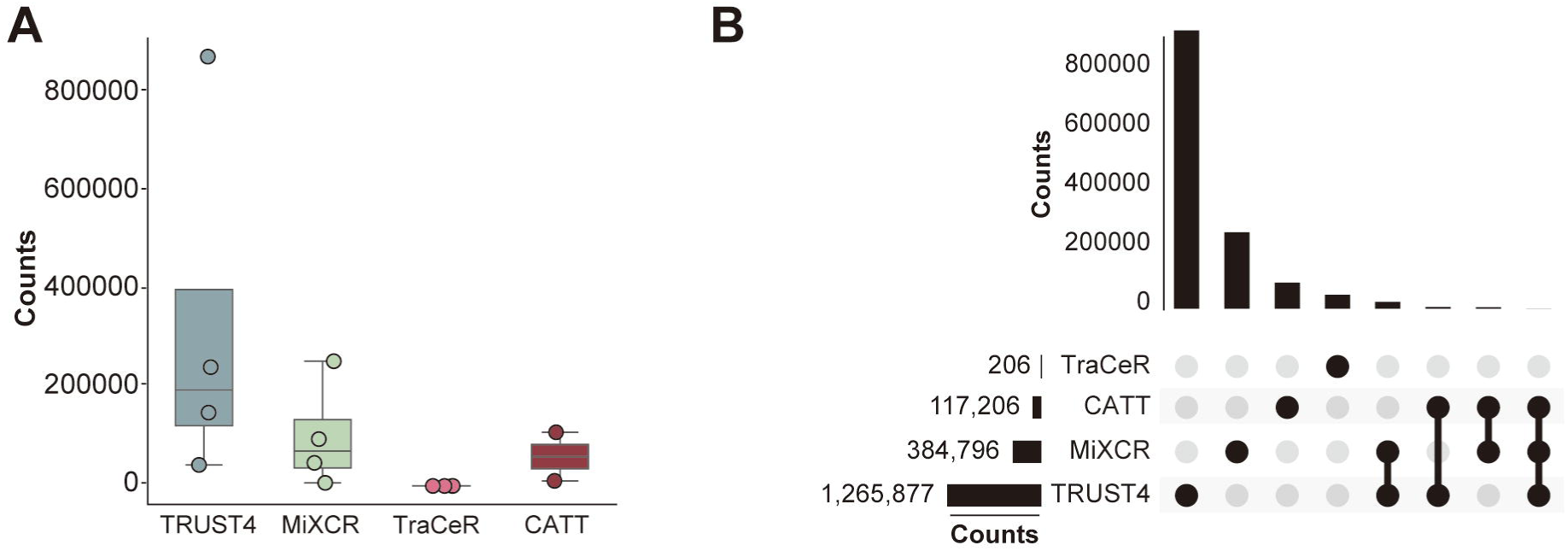
Comparative analysis of CDR3β construction methods on bulk TCR-seq data. **A.** boxplot represents the number of unique CDR3β sequences assembled by each method. **B.** Upset plots represent the overlap of unique CDR3β sequences assembled by each method.

**Supplementary Figure 10.**
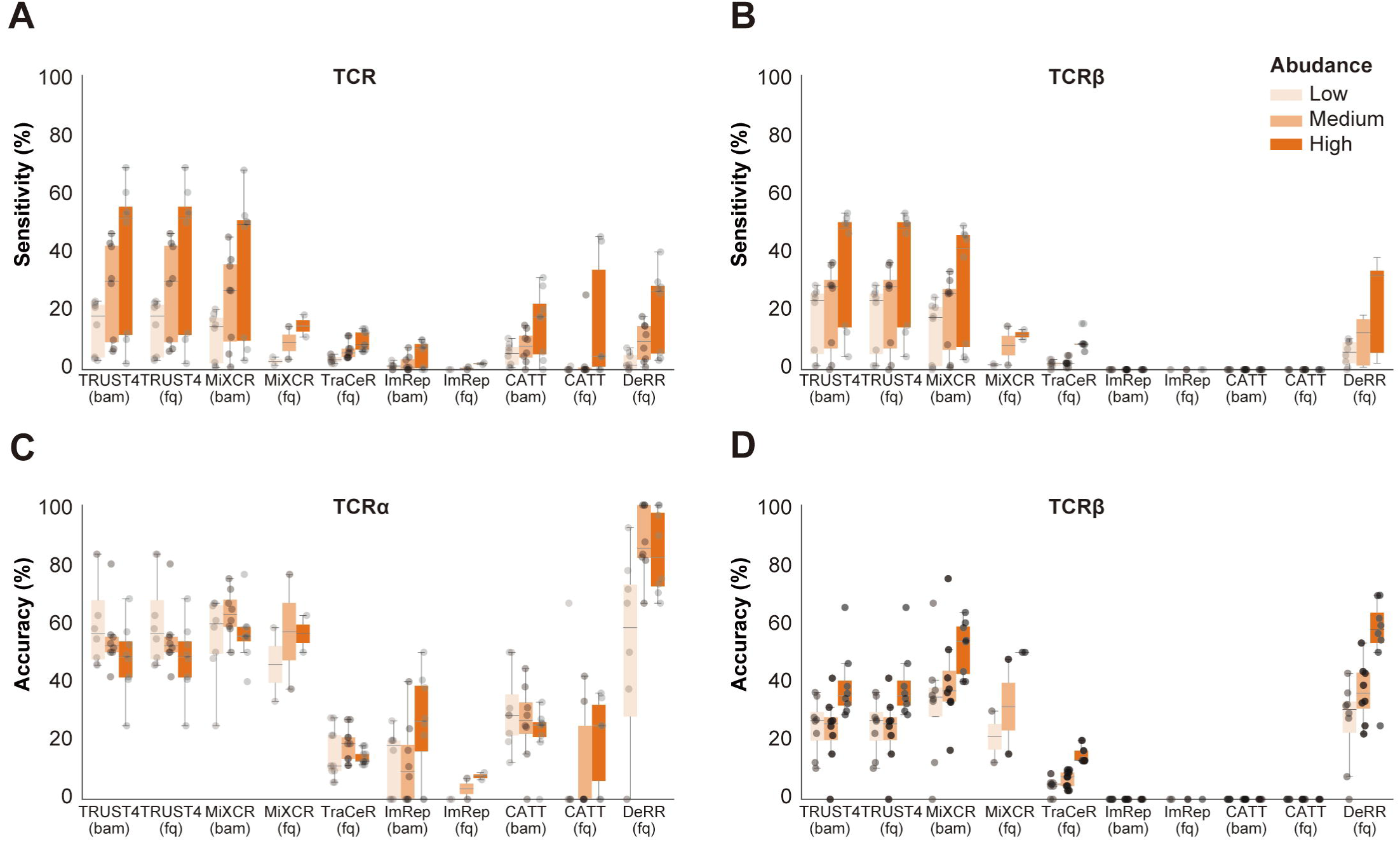
Performance assessment of methods on pseudo bulk RNA-seq with varying TCR abundance. A-D. Box plots represent the sensitivity **(A,B)** and accuracy **(C,D)** of TCRα and TCRβ assembled by different methods at varying TCR abundance. The line in the middle represents the median; boxes represent the 25th (bottom) and 75th (top) percentiles; and whiskers represent the minimum and maximum points within 1.5 times the interquartile range. fq, FASTQ format; bam, Binary Alignment Map format.

